# Spatial transcriptomics reveals tumour cell-intrinsic and microenvironmental features associated with survival in high-grade serous ovarian carcinoma

**DOI:** 10.64898/2026.10.02.755833

**Authors:** Kamran Khan, Weitao Lin, Elena Denisenko, Rui Hou, Sanela Bilic, Yu Yu, Yin Huang, Peiwen Liu, Paul A. Cohen, Alistair R. R. Forrest

## Abstract

High-grade serous ovarian carcinoma (HGSOC) is the most common form of ovarian carcinoma and accounts for the majority of deaths from the disease, with fewer than half of patients surviving five years after diagnosis. Bulk transcriptomic profiling has identified molecular subtypes and gene-expression signatures associated with shorter or longer survival. However, because bulk profiling measures a composite of tumour and microenvironmental gene expression, the cellular origins of these signatures and their relationship to the tumour microenvironment remain unresolved. To address this, we applied Xenium spatial transcriptomics to treatment-naïve HGSOC tumours from 122 patients using a custom 480-gene panel designed to measure tumour-intrinsic gene expression and distinguish immune and stromal cell populations in the surrounding microenvironment. Analysis of tumour-intrinsic gene expression identified 61 genes nominally associated with overall survival, from which Elastic-Net modelling prioritised five genes associated with longer survival and nine associated with shorter survival. Notably, 46 of the 61 associations did not reach the nominal significance threshold when expression was instead aggregated across all cells from the same patients, highlighting how whole-tissue expression can obscure tumour-cell-specific survival associations. Complementary analysis of tumour-proximal cell populations revealed that multiple immune populations were associated with longer survival, whereas two fibroblast populations were associated with shorter survival. Integrating these analyses revealed that survival-associated tumour-intrinsic gene expression was correlated with the abundance of neighbouring immune and stromal populations that were themselves associated with survival. Pairwise Cox regression demonstrated substantial overlap in the prognostic information captured by these features, while analysis of individual gene–cell combinations showed that survival associations differed according to the abundance of neighbouring populations. Consensus clustering further identified four patient groups with distinct combinations of tumour-intrinsic expression and microenvironmental composition, broadly separating patients with longer and shorter survival. Together, this cohort-scale spatial analysis identifies tumour-intrinsic and microenvironmental features associated with HGSOC survival and reveals how their correlated patterns and combined associations relate to patient outcome.

## Introduction

High-grade serous ovarian carcinoma (HGSOC) exhibits substantial variation in patient outcomes, and bulk transcriptomic analyses have classified HGSOC into four gene-expression subtypes (mesenchymal, proliferative, immunoreactive and differentiated), each associated with differences in survival^1^. These signatures were later refined and translated into a targeted 55 gene diagnostic panel for subtype prediction^2^. However, bulk RNA profiling captures a mixture of tumour and stromal signals. For example, the immunoreactive subtype largely reflects immune infiltration, whereas the proliferative and differentiated subtypes primarily capture tumour-intrinsic gene-expression patterns. The mixed cellular origins of these signatures complicate their biological interpretation and may contribute to differences in subtype classification across studies^3,4^. Notably, different regions within the same tumour can correspond to different transcriptional subtypes, highlighting a limitation of bulk tumour profiling^5^.

Independent studies using immunohistochemistry and immunofluorescence have demonstrated that immune infiltration within the tumour microenvironment is strongly associated with survival in HGSOC. Increased numbers of CD8+ T cells correlate with longer survival^6,7^, while tumour-infiltrating B cells and IgA+ plasma cells are also associated with favourable outcomes^8^. More recently, spatial proteomic approaches have further resolved immune and stromal microenvironments linked to prognosis. Imaging mass cytometry identified tertiary lymphoid structures associated with longer survival and fibroblast-rich microenvironments associated with shorter survival^9^. Similarly, multiplex immunofluorescence analyses have reported associations between cytotoxic CD8+ T cells, CD66+ cells, and regulatory T cells in proximity to tumour cells and longer overall survival^10^. Together, these studies demonstrate that the spatial organisation of immune and stromal populations is associated with survival in HGSOC, but do not establish how these microenvironmental features relate to survival-associated gene expression within neighbouring tumour cells.

In addition to transcriptional and immune features, genomic alterations such as *CCNE1* amplification and deletion of chromosome region 6q24.2–6q26 have been associated with differences in HGSOC survival^11,12^. Genomic heterogeneity may also contribute to differences in the local tumour microenvironment. Our previous spatial transcriptomic analysis identified genetically distinct tumour subclones within individual HGSOC tumours that expressed different ligand repertoires and were associated with distinct neighbouring immune and stromal populations^5^. These observations raise the possibility that tumour-intrinsic gene-expression differences contribute to variation in microenvironmental composition, providing a rationale for examining these features together in relation to patient outcome.

To investigate these relationships more directly, we developed a custom 480-gene panel and applied it to treatment-naïve HGSOC tissue microarrays (TMAs) linked to overall survival data. The panel comprised 260 genes selected to capture tumour-intrinsic gene expression and 220 genes identifying infiltrating immune and stromal cell populations.

Using this spatially resolved dataset, we addressed three questions. First, which tumour-intrinsic genes are associated with overall survival? Second, which tumour-proximal microenvironmental cell populations are associated with overall survival? Third, are tumour-intrinsic gene-expression patterns associated with particular tumour-proximal microenvironmental compositions, and do these features provide overlapping or independent prognostic information, and can their combined profiles distinguish patient groups with different survival outcomes?

## Results

### Spatial profiling of HGSOC using a custom 480-gene Xenium panel

To systematically investigate associations between tumour-intrinsic gene expression, the tumour-proximal microenvironment and overall survival in HGSOC, we profiled tumour cores from 122 treatment-naïve patients using Xenium *in situ* spatial transcriptomics (**Figure 1A**). Median overall survival (OS) was 33 months (IQR 14.7–47.1) and median age at diagnosis was 65 years (IQR 56–74) (**Supplementary Table S1**).

**Figure 1.**
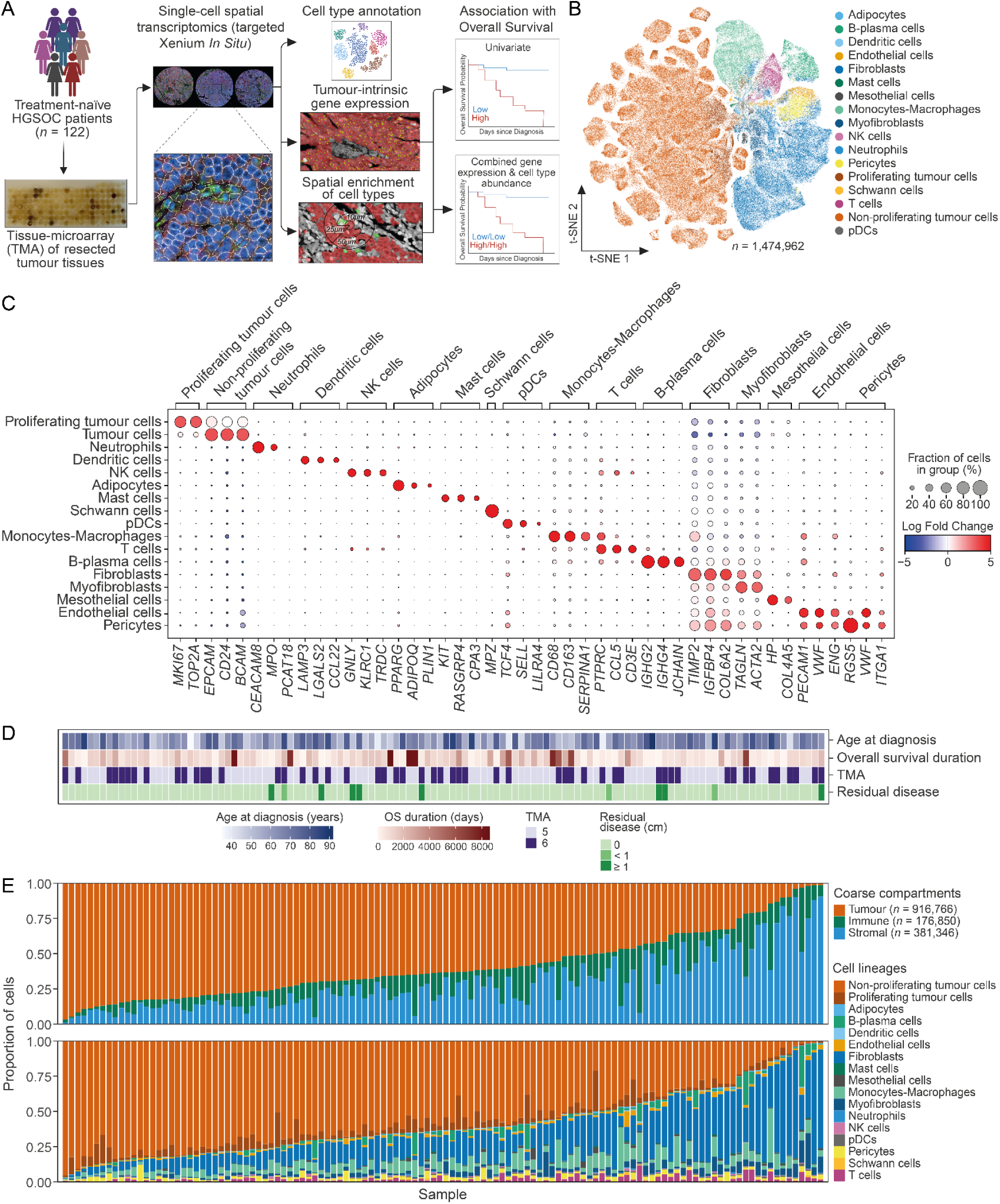
Spatial transcriptomic profiling and cellular composition of the HGSOC cohort. **A.** Schematic workflow of the experimental pipeline. Resected, treatment-naïve tumour samples from a cohort of 122 patients with high-grade serous ovarian carcinoma (HGSOC) were assembled into tissue microarrays (TMAs) and profiled using Xenium in situ spatial transcriptomics with a custom 480-gene panel. Downstream workflows encompass cell type annotation, cell-intrinsic gene expression profiling, spatial architectural analysis, and clinical correlation modelling with overall survival. **B.** t-SNE projection of 1.47 million high-quality single cells, colour-coded by 17 broad cell lineages spanning tumour, stromal, and immune compartments. **C.** Dot plot representing the canonical marker genes expressed by the 17 annotated lineages. The size of each dot represents the percentage of cells within a lineage expressing a given marker, and the colour intensity reflects the log-fold change of expression compared to all other cell populations. **D.** Heatmap illustrating the baseline clinicopathological characteristics of the 122-patient cohort, including overall survival duration (days), age at diagnosis, TMA assignment, and residual disease status. Samples are arranged in an identical left-to-right patient order as shown in panel E to facilitate direct visual comparison. **E.** Stacked bar plots displaying the proportional cellular distributions across individual core samples, maintained in the same patient alignment as panel D. The upper panel depicts the distribution of coarse compartments (Tumour, Immune, and Stromal); the lower panel displays the detailed composition of all 17 fine-grained lineages across the samples. Figure 1A template was created in BioRender. Khan, M.A.A.K. (2026) https://BioRender.com/yzi73t2

To resolve tumour-intrinsic gene expression and the cellular composition of the microenvironment, we developed a custom 480-gene Xenium panel by integrating and prioritising genes from 14 major HGSOC datasets^2,4,13–23^. Panel design balanced 220 markers selected to distinguish immune and stromal cell populations with 260 genes selected to interrogate tumour-intrinsic biology, including HGSOC molecular subtype and survival-associated genes, genes within recurrent copy-number alterations, mismatch repair genes, candidate circulating protein biomarkers and genes identified in our previous spatial transcriptomic study^5^ (**Supplementary Table S2, Methods**). The panel was applied to two tissue microarrays (TMAs), comprising 1-mm tumour cores (**Methods**). Following transcript reassignment and quality control filtering, 208 tissue cores from 122 patients were retained for downstream analysis. In total, we mapped 1,474,962 high-quality single cells, yielding approximately 12,000 cells per patient.

Supervised marker-guided cell type annotation (**Methods**) partitioned the single-cell dataset into 17 broad cell lineages spanning the malignant, stromal, and immune compartments, recovering 916,766 tumour cells, 381,346 stromal cells, and 176,850 immune cells (**Figure 1B–C**). To visualise the overall cellular landscape, all 1.47 million high-quality cells were projected into a two-dimensional t-SNE embedding, revealing distinct clusters corresponding to the 17 annotated cell lineages (**Figure 1B**). These annotations were supported by lineage-specific expression of canonical marker genes (**Figure 1C**). Tumour cells were identified by expression of *EPCAM*, *CD24* and *BCAM*, with proliferating tumour cells distinguished by expression of *MKI67* and *TOP2A*. Immune cell populations were resolved by lineage-specific markers, including *CEACAM8* and *MPO* (neutrophils), *LAMP3* (dendritic cells), *GNLY* and *KLRC1* (NK cells), *TCF4* (plasmacytoid dendritic cells, pDCs), *CD68* and *CD163* (monocytes-macrophages), *PTPRC* and *CD3E* (T cells), *IGHG2, IGHG4* and *JCHAIN* (B-plasma cells), and *CPA3* (mast cells). Similarly, structural and stromal lineages displayed high marker specificity, including *PPARG* (adipocytes), *MPZ* (Schwann cells), *TIMP2* and *IGFBP4* (fibroblasts), *TAGLN* and *ACTA2* (myofibroblasts), *HP* (mesothelial cells), *PECAM1* and *ENG* (endothelial cells), and *RGS5* (pericytes).

Profiling cellular composition across the cohort revealed substantial inter-patient heterogeneity. Tumour cells constituted a median of 63.4% of all cells (IQR 46.4%–76.8%), followed by stromal cells (22.2%, IQR 13.8%–38.4%), and immune cells (9.4%, IQR 4.9%–15.7%). Within the stromal compartment, fibroblasts were the dominant lineage, followed by myofibroblasts, whereas the remaining structural populations were comparatively sparse. Monocytes-macrophages represented the most abundant immune population, followed by B-plasma cells and T cells (**Figure S1B**). Cell-type composition varied substantially between patients and was not obviously explained by baseline clinicopathological variables (**Figure 1D–E**).

Because the two TMAs were processed across multiple Xenium slides (TMA5 across three slides and TMA6 across two slides), we next assessed technical reproducibility. Linear modelling of median cell-type abundances across the 17 major cell showed strong concordance between independent Xenium runs (**Supplementary Table S3**) and between the two TMAs (R² = 0.96, p = 7.3 × 10⁻¹²), indicating minimal batch effects (**Figure S1C**). Cells from different Xenium slides were also well mixed within the t-SNE embedding (**Figure S1D**), indicating limited technical variation in cell-type composition. Tumour cells nevertheless showed distinct patient-specific expression patterns, whereas stromal and immune cells from different patients occupied largely overlapping regions of the embedding.

Although the broad lineage classifications captured the major cellular components of HGSOC, several of these compartments are known to contain functionally distinct cell states. To resolve cellular heterogeneity within these broad lineages, we further subdivided lymphoid, myeloid and structural compartments into finer cellular states (**Figure S1E**), defining a total of 51 distinct cell populations across the cohort. Within the B-cell compartment, *IGHG+* B-plasma cells were the most abundant subset, while *CCL5+* T cells represented the most abundant T-cell subpopulation. Similarly, *GLUL+* and *CD163+* macrophages, *EGR1+* and *IGFBP5+* fibroblasts, and venous endothelial cells constituted the most abundant populations within their respective compartments (**Figure S1F**).

To determine whether baseline clinical variables including age and residual disease were independently associated with survival, clinicopathological variables together with sample-level tumour cell abundance were evaluated using multivariable Cox proportional hazards modelling. Increasing age at diagnosis was associated with shorter survival (hazard ratio (HR) = 1.02, 95% CI: 1.00–1.04, p = 0.02), whereas no other clinicopathological variables, including tumour cell abundance, were significantly associated with overall survival (**Figure S1A**). We next investigated tumour-intrinsic gene expression and tumour-proximal microenvironmental features associated with survival.

### Tumour-intrinsic gene expression identifies survival-associated features in HGSOC

Expression of numerous genes in HGSOC has been associated with variable clinical outcomes^1,23–26^. However, the vast majority of these reported survival-associated genes were identified using bulk tissue transcriptomics, in which RNA derived from malignant, stromal and immune cells is measured as a single mixture, obscuring the cellular origin of individual transcripts.

To identify tumour-intrinsic gene expression associated with survival, we restricted our downstream analysis to cells annotated as tumour cells, including both proliferating and non-proliferating tumour cells (**Figure 2A**). For each patient, single-cell expression profiles from all tumour cells were aggregated to generate a patient-level pseudo-bulk expression profile. Hereafter, we refer to gene expression measured specifically within the tumour-cell compartment as tumour-intrinsic gene expression. Initial assessment of these pseudo-bulk profiles by two-dimensional principal component analysis (PCA) revealed a clear separation by TMA runs, indicating a prominent technical batch effect. We therefore performed batch-correction with ComBat^27^ to reduce this technical variation (**Methods**). Following correction, run-specific clustering was no longer evident (**Figure S2A**). Corrected median transcript abundances were also highly correlated between the TMA runs (R^2^ = 0.99, p < 2.2 x 10^-16^; **Figure S2B**).

**Figure 2.**
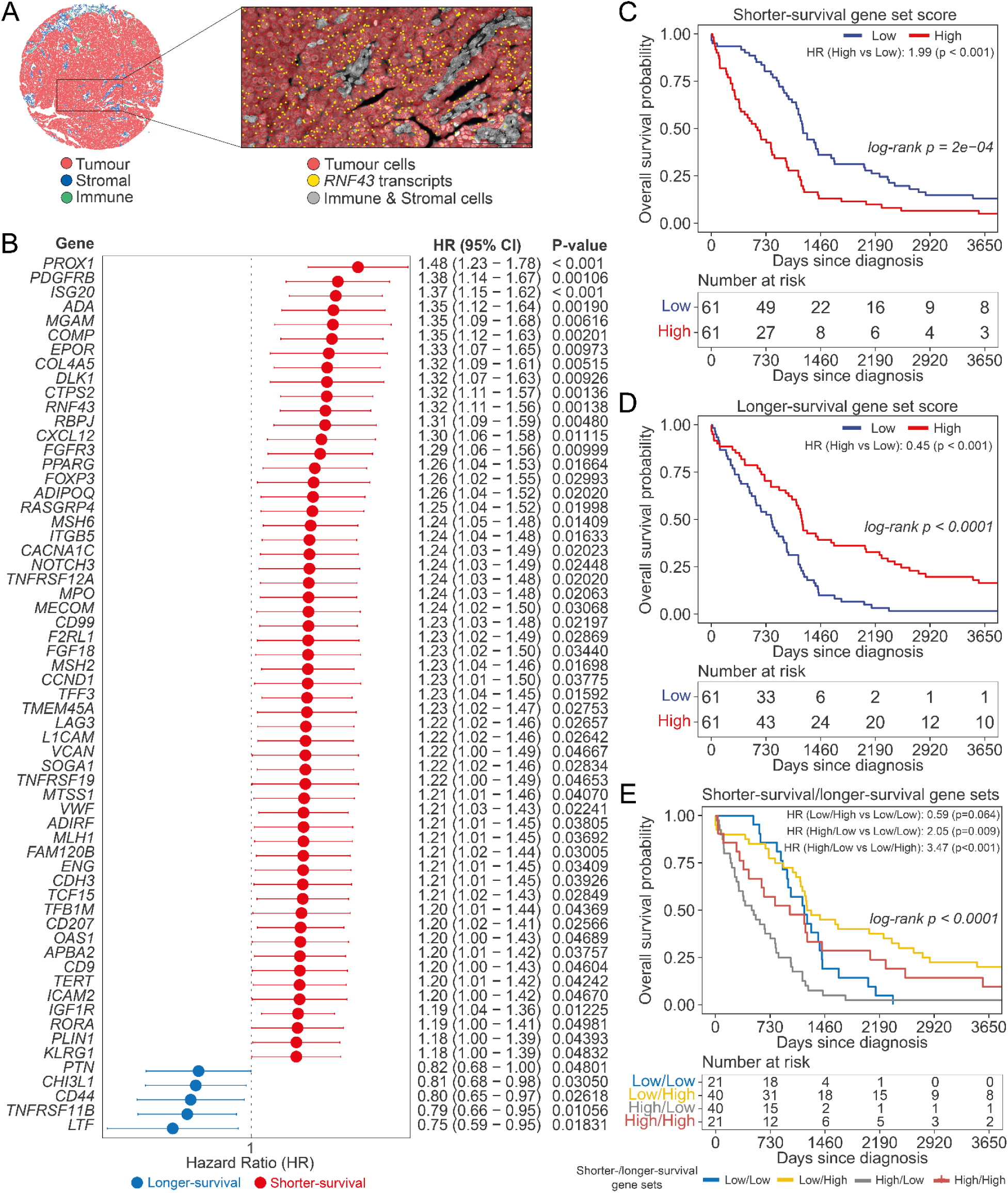
**Tumour**–**cell-intrinsic gene expression patterns are associated with HGSOC clinical outcomes. A.** Representative Xenium TMA core and zoomed-in region showing spatial localisation of *RNF43* transcripts (yellow dots) within tumour cell segments (red), distinct from stromal and immune segments (grey). **B.** Forest plot illustrating univariate Cox proportional hazard ratios (HR) of tumour-intrinsic genes associated with survival, utilising gene expression as continuous variables. Data are presented as HRs with 95% confidence intervals (CI) and corresponding nominal p-values. Genes with longer-survival associations (HR < 1) are shown in blue, and genes with shorter-survival associations (HR > 1) are shown in red. **C.** Kaplan–Meier curves for the shorter-survival gene set score (9 Elastic–net–prioritised shorter-survival genes), with patients stratified into high and low groups using a median split. **D.** Kaplan–Meier curves for the longer-survival gene set score (5 Elastic–net–prioritised longer-survival genes), with patients stratified into high and low groups using a median split. **E.** Combined analysis of shorter- and longer-survival gene sets, stratifying patients into four groups based on high/low gene set scores. All Kaplan–Meier curves display only the first 10 years of overall survival for visualisation purposes; all analyses used the full survival duration. Significance metrics for the Cox proportional hazard models were derived using the Wald statistic (*p < 0.05, **p < 0.01, ***p < 0.001). Global p-values for Kaplan–Meier survival curves were determined using the non-parametric log-rank test, which compares the survival distributions across patient groups over the observed follow-up period. All reported p-values are nominal and were not adjusted for multiple testing.

The batch-adjusted pseudo-bulk expression matrices were z-score scaled across the cohort and evaluated using univariate Cox proportional hazards regression models **(Methods)**. This analysis identified 61 genes whose tumour-intrinsic expression was nominally associated with overall survival (p < 0.05), including 56 associated with shorter survival and five associated with longer survival (**Figure 2B**). One gene remained significantly associated with overall survival after Benjamini-Hochberg correction for multiple testing across the targeted gene panel (FDR < 0.05) (**Supplementary Table S4**). The 61 nominally associated genes were subsequently used as a candidate set for multivariable feature selection.

To determine how restricting expression measurements to tumour cells affected the identification of survival-associated genes, we repeated the analysis using whole-tissue pseudo-bulk profiles generated by aggregating expression across all cells from each patient. Of the 61 genes associated with overall survival using tumour-cell pseudo-bulk expression, 15 were also associated with survival using whole-tissue pseudo-bulk expression, whereas 46 were identified only in the tumour-cell analysis (**Figure S3A-B**). Conversely, 21 genes were associated with survival only using whole-tissue pseudo-bulk expression. Examination of their expression across the annotated cell populations showed that these whole-tissue-specific associations predominantly reflected expression by non-malignant populations: four genes were specific to *IGFBP5+* fibroblasts, 11 were broadly expressed across fibroblast populations, two were expressed by macrophages, two by myofibroblasts and one by plasmacytoid dendritic cells, while the remaining gene was *IGHM*, expressed by the *IGHM+* B-plasma-cell population (**Figure S3C; Supplementary Table S5**). Thus, restricting expression measurements to tumour cells identified 46 survival associations that did not reach the nominal significance threshold in whole-tissue pseudo-bulk analysis, while genes identified only using whole-tissue expression predominantly reflected expression from non-malignant cell populations.

Several of the survival-associated genes mapped to recurrently amplified genomic regions in HGSOC. Five genes associated with shorter survival (*NOTCH3*, *ADIPOQ*, *RASGRP4*, *MECOM* and *MTSS1*) and one associated with longer survival (*TNFRSF11B*) resided within chromosomal regions amplified in more than 10% of TCGA HGSOC samples, indicating that these expression associations occur within genomic regions recurrently altered across patients. Several survival-associated genes are also clustered within the same or neighbouring genomic regions. *CD9*, *KLRG1*, *LAG3* and *VWF* were located within a recurrently amplified region at 12p13.31, while three additional gene pairs, *MECOM* and *ADIPOQ*, *TNFRSF11B* and *MTSS1*, and *NOTCH3* and *EPOR*, were located in close genomic proximity. Thus, the proximity of some survival-associated genes within recurrently amplified regions raises the possibility that their correlated expression reflects shared copy-number alterations rather than independent regulation (**Supplementary Table S6**).

Because several of the survival-associated genes are more commonly recognised as markers of non-malignant cell populations, we sought to confirm that the expression attributed to tumour cells represented genuine tumour-cell expression. These included *FOXP3* and *PROX1*, canonical markers of regulatory T-cells and lymphatic endothelial cells respectively. Examination of the 61 survival-associated genes across the Xenium dataset showed that their expression within tumour cells varied substantially between patients and that several were also expressed by non-malignant populations (**Figure S4-5**). Direct inspection confirmed transcripts from all 61 genes within annotated tumour-cell segments (**Figure S6**). Human Protein Atlas immunohistochemistry showed moderate-to-strong tumour-cell protein staining for 34 genes in serous ovarian carcinoma (**Figure S7, Supplementary Table S4**), while independent single-cell RNA-seq datasets detected all 61 genes in tumour cells in at least a subset of HGSOC samples (**Figure S8, Supplementary Table S4**). Together, these orthogonal lines of evidence support expression of these genes within malignant cells, including genes more commonly used as markers of other cell types.

As many of the 61 candidate genes exhibited coordinated co-expression patterns (**Figure S9**), we reasoned that some represented correlated features carrying redundant survival-associated information. We therefore applied Elastic-Net regularised regression modelling with a λ-1se penalty to prioritise a smaller set of survival-associated genes. This reduced the 56 shorter-survival genes to nine prioritised features, including *PROX1*, *L1CAM*, *CTPS2*, *ISG20*, *COMP*, *ADA*, *COL4A5*, *CCND1*, and *RNF43*, while retaining all five longer-survival genes (**Figure S10A–B; Supplementary Table S7**). Pairwise correlation analysis of the 14 prioritised genes showed limited shared variance, and most associations were retained when the prioritised genes were evaluated jointly (**Figure S10C–D**).

We next evaluated the collective survival associations of these genes by calculating ssGSEA scores independently for the nine–gene shorter-survival and five–gene longer-survival gene sets (**Methods**). Median–based patient stratification using these scores revealed marked survival differences for both the shorter-survival gene set (**Figure 2C**; HR = 1.99, log–rank p = 2 × 10⁻⁴) and the longer-survival gene set (**Figure 2D**; HR = 0.45, log–rank p < 0.0001). Joint analysis of both scores revealed a gradient of survival outcomes, with patients exhibiting high shorter-survival and low longer-survival gene set scores experiencing the shortest survival, and patients exhibiting low shorter-survival and high longer-survival gene set scores experiencing the longest survival (**Figure 2E**; global log–rank p < 0.0001). In a Cox model containing both scores, each retained an association with overall survival after mutual adjustment (HR_shorter-survival_ = 1.45, 95% CI: 1.17–1.81, p < 0.001; HR_longer-survival_ = 0.64, 95% CI: 0.52–0.78, p < 0.001), indicating that the two gene sets captured complementary survival-associated information (**Figure S10E**).

Finally, to summarise the combined survival-associated information captured by the prioritised tumour–intrinsic gene sets, we integrated the shorter– and longer-survival signatures into a continuous composite score (**Methods**). Median stratification separated patients into groups with distinct survival trajectories (**Figure S11A**), with patients in the higher–score group exhibiting shorter overall survival than those in the lower–score group (HR = 3.30, 95% CI: 2.21–4.93, log–rank p < 0.0001). Time-dependent receiver operating characteristic analysis further evaluated discrimination across multiple follow-up intervals (**Figure S11B**). To assess whether the survival association was retained in patients not used for feature selection or model fitting, we performed nested 10-fold cross-validation (**Methods**). In each training fold, survival-associated genes were identified and Elastic-Net regularisation was applied to prioritise features, with the resulting model used to generate scores for the held-out patients. Each patient’s out-of-fold score was therefore generated using a model developed without that patient’s data. The resulting out-of-fold scores remained associated with overall survival (**Figure S11C**; HR = 1.92, 95% CI: 1.03–3.55, p = 0.0393). As the composite score was derived and internally evaluated within the same cohort, its performance will require validation in an independent cohort.

Together, these analyses identify a broad set of tumour-intrinsic expression features associated with overall survival in HGSOC, prioritise a smaller set with limited expression redundancy, and show that their combined expression captures complementary survival-associated information.

### Tumour–proximal immune and fibroblast populations show opposing associations with overall survival

To quantify the cellular composition immediately surrounding tumour cells, we measured non-malignant cell abundance within defined distances of individual tumour cells rather than across the entire tissue core. Using Squidpy’s^28^ spatial neighbourhood graphs, we quantified neighbouring non-malignant cells within incremental radii of 10, 25, 50, 100, 250 and 500 μm around each tumour cell (**Methods; Figure 3A**). This enabled systematic quantification of tumour-proximal immune and stromal cell abundance across a range of neighbourhood sizes (**Figure S12A**). We first evaluated tumour-proximal cell abundance as a continuous variable using independent univariate Cox proportional hazards regression models. *IGFBP5+* fibroblast abundance was consistently associated with shorter overall survival across all evaluated radii, whereas *IGHM+* B-plasma cells were associated with longer overall survival irrespective of distance from tumour cells (**Figure S12B**). Additional immune populations demonstrated distance-dependent associations with survival. *IL32+* T cells and *CXCL9+* macrophages were associated with longer survival only within the immediate 10 μm neighbourhood, and regulatory T cells at 10-50 μm, whereas dendritic cells were associated with longer survival only at more distant neighbourhoods (250-500 μm) (**Figure S12B**). When cell abundance across the entire tissue core was analysed, the associations of *IGHM+* B-plasma cells with longer survival and *EGR1+* and *IGFBP5+* fibroblast abundance with shorter survival remained significant, while *CD163+* macrophages were associated with shorter survival (**Figure S13**). All other associations were detected only in the tumour-proximal neighbourhood analyses.

**Figure 3.**
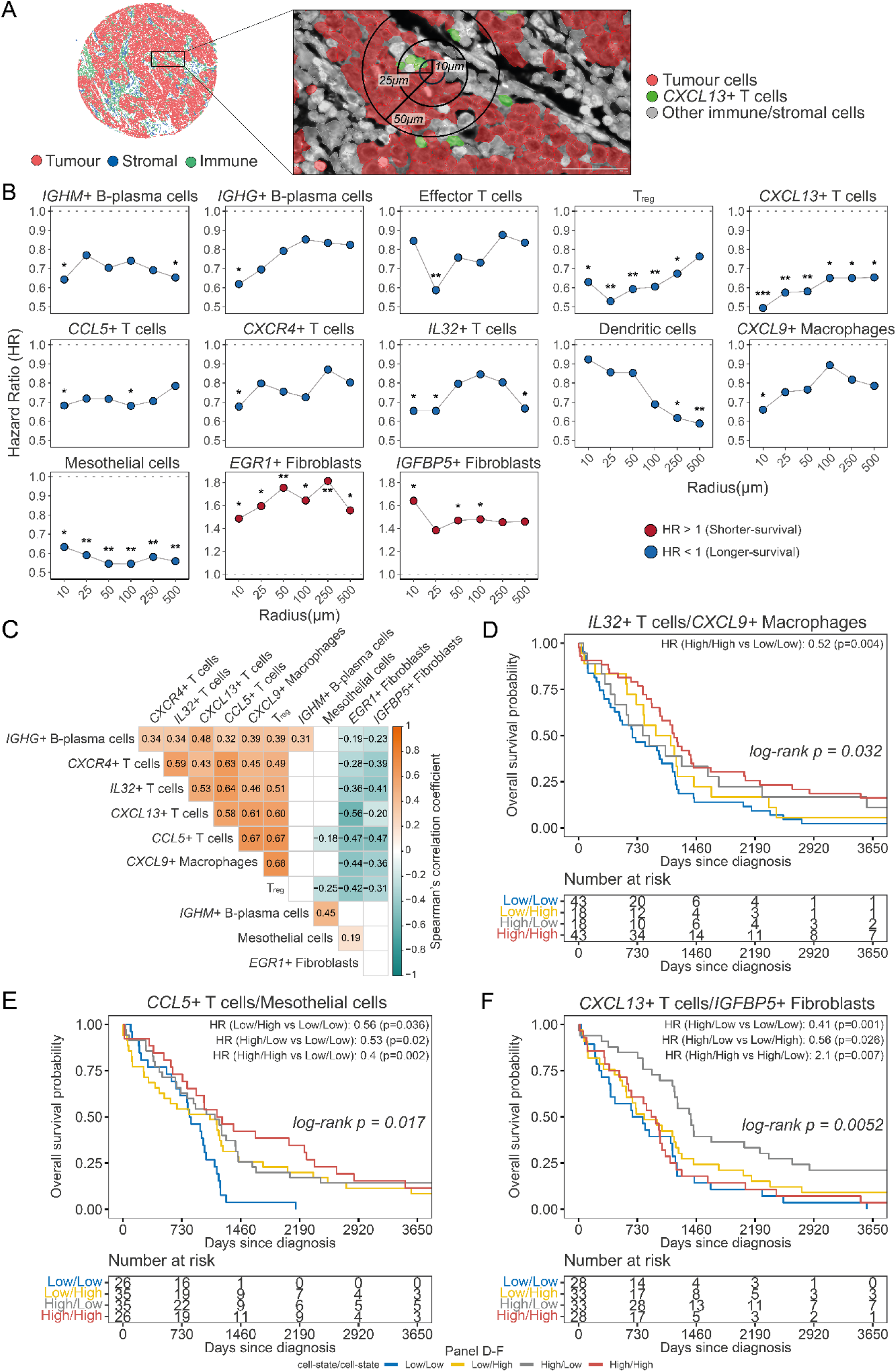
Spatial microenvironmental features surrounding tumour cells are associated with overall survival in HGSOC. **A.** Spatial mapping of tumour, stromal and immune cell segments, illustrating the quantification of non-malignant cell populations within concentric radii surrounding tumour cells. **B.** Hazard ratios for associations between tumour-proximal microenvironmental cell states and overall survival across spatial radii (10–500 μm). Each curve represents a microenvironmental cell state, with hazard ratios plotted as a function of distance from tumour cells. Increasing radii capture progressively broader spatial contexts, from local cellular neighbourhoods to broader tumour ecosystem organisation. High- and low-abundance groups for the underlying categorical Cox proportional hazards analyses were defined using 75th and 25th percentile thresholds, respectively. **C.** Hierarchically clustered Spearman correlation matrix showing pairwise relationships between microenvironmental cell states identified as outcome-associated at the 10 μm radius. Only statistically significant pairwise correlations (p < 0.05) are shown. **D.** Kaplan–Meier curves comparing patient groups jointly stratified by *IL32+* T-cell and *CXCL9+* macrophage abundance within 10 μm of tumour cells, illustrating the survival association of concurrent enrichment of populations within the coordinated longer-survival-associated immune programme. **E.** Kaplan–Meier curves comparing patient groups jointly stratified by *CCL5+* T-cell and mesothelial-cell abundance within 10 μm of tumour cells, illustrating complementary longer-survival-associated effects of populations from relatively independent prognostic axes. **F.** Kaplan–Meier curves comparing patient groups jointly stratified by *CXCL13+* T-cell and *IGFBP5+* fibroblast abundance within 10 μm of tumour cells, illustrating opposing associations of immune and stromal populations with overall survival. Panels D–F use median splits to define high- and low-abundance groups. Kaplan–Meier curves display only the first 10 years of overall survival for visualisation purposes; all statistical analyses were performed using the full survival duration. For B, significance metrics for Cox proportional hazards models were derived using the Wald statistic (*p < 0.05, **p < 0.01, ***p < 0.001). Global p-values for Kaplan–Meier survival curves were determined using the non-parametric log-rank test, which compares the survival distributions across patient groups over the observed follow-up period. For D–F, only significant contrasts (p < 0.05) are shown. All reported p-values are nominal and were not adjusted for multiple testing.

Because the abundance distributions of many cell populations were highly skewed across the cohort, we next compared patients with high (75^th^ percentile) and low (25^th^ percentile) levels of infiltration using categorical Cox regression models (**Methods**). This analytical framework revealed associations of multiple immune and stromal cell subsets with overall survival (**Figure 3B, Figure S12C**),recapitulating the continuous trends described above while identifying additional survival-associated populations. Using this approach, we found that the associations between *CXCL13+* T cells and mesothelial cells and longer survival, together with the association between *EGR1+* fibroblasts and shorter survival, were preserved across all evaluated radii (**Figure 3B**). Other populations displayed tightly distance-specific associations, including *IGHM+* B-plasma cells (10 and 500 μm), *IGHG+* B-plasma cells (10 μm), effector T-cells (25 μm), *CXCR4+* T-cells (10 μm), *CXCL9+* Macrophages (10 μm), *CCL5+* T-cells (10 and 100 μm), and *IL32+* T-cells (10, 25, and 500 μm). Notably, regulatory T-cells were associated with longer survival across almost all the evaluated distances (10, 25, 50, 100, and 250 μm). In contrast, *IGFBP5+* fibroblasts were associated with shorter overall survival only when located close to tumour cells (10, 50, and 100 μm) (**Figure 3B**). Among immune populations associated with longer survival, associations were generally strongest close to tumour cells and weakened with increasing distance. In contrast, dendritic cells displayed the opposite spatial trend: their abundance was associated with longer survival only at more distal distances from tumour cells (**Figure 3B**).

We next examined whether these survival-associated cell populations exhibited coordinated abundance patterns. Pairwise Spearman correlations across all spatial distances revealed significant correlations among multiple survival-associated populations (**Figure 3C, Figure S14**). The strongest positive correlations were between regulatory T cells and *CXCL9+* macrophages (*r* = 0.68), followed by *CCL5+* T cells and *CXCL9+* macrophages (*r* = 0.67), and *CCL5+* and regulatory T cells (*r* = 0.67). These populations formed part of a larger cluster of positively correlated, longer-survival-associated immune populations, comprising regulatory, *CCL5+*, *CXCR4+*, *IL32+* and *CXCL13+* T cells, *CXCL9+* macrophages and *IGHG+* B-plasma cells, which we refer to as the T-cell-enriched immune cluster. A second, distinct longer-survival-associated cluster comprised mesothelial cells and *IGHM+* B-plasma cells, whose abundances were also positively correlated (*r* = 0.45).

Populations within these longer-survival-associated clusters were negatively correlated with the two fibroblast populations associated with shorter survival, with the strongest negative correlation observed between *CXCL13+* T cells and *EGR1+* fibroblasts (*r* = −0.56). In contrast, we observed no significant correlation between the abundances of *IGFBP5+* and *EGR1+* fibroblasts (**Figure 3C, Figure S14**).

To assess the extent to which survival associations were shared among correlated cell populations, we next performed pairwise joint Cox models in which each survival-associated population was evaluated after accounting for a second population. Within the T-cell-enriched immune cluster, associations for several populations were attenuated when modelled alongside other members of the cluster, consistent with substantial overlap in their survival-associated information (**Figure S15, Supplementary Table S8**). In contrast, *IGHM+* B-plasma cells and mesothelial cells retained their associations with longer survival after accounting for populations from the T-cell-enriched immune cluster. However, neither population retained a significant association after accounting for the other. Together with their positively correlated abundances, these findings suggest that *IGHM+* B-plasma cells and mesothelial cells capture overlapping survival-associated information that is relatively independent of the broader T-cell-enriched immune cluster (**Figure 3C, Figure S14, Supplementary Table S8**).

A similar distinction was observed between the two shorter-survival-associated fibroblast populations. The association of *IGFBP5+* fibroblasts with shorter survival was retained after adjustment for *EGR1+* fibroblasts and multiple other neighbouring cell populations, whereas the association of *EGR1+* fibroblasts was attenuated after accounting for *IGFBP5+* fibroblasts. Thus, despite the absence of a significant correlation between their abundances, *IGFBP5+* fibroblasts captured a larger component of the prognostic information associated with these fibroblast populations, with *EGR1+* fibroblasts providing less independent prognostic information (**Figure S15, Supplementary Table S8**). We next examined how these patterns were reflected in patient-level survival.

Within the T-cell-enriched immune cluster, *CXCL9+* macrophages and *IL32+* T cells provided a representative example of the shared prognostic signal identified by the correlation and joint Cox analyses. Patients with high abundance of both populations had significantly longer overall survival than those with low abundance of both populations (HR = 0.52, p = 0.004), while patients with high abundance of either population alone showed intermediate survival (**Figure 3D**). Pairwise comparisons involving these intermediate groups were not significant (**Supplementary Table S8**). Thus, the longest survival was observed when both correlated immune populations were abundant.

Populations representing distinct longer-survival-associated patterns showed a different relationship. *CCL5+* T cells and mesothelial cells belonged to separate abundance clusters and retained relatively independent prognostic associations in joint Cox models (**Figure 3C, Figure S15**). Patients with high abundance of either *CCL5+* T cells or mesothelial cells, or both populations, had significantly longer overall survival than patients with low abundance of both (**Figure 3E**). Although patients with high abundance of both populations showed a trend towards more favourable survival than those with high abundance of either population alone, these differences were not significant (**Supplementary Table S9**). These findings are consistent with *CCL5+* T cells and mesothelial cells representing distinct favourable microenvironmental features, either of which is associated with improved survival.

Finally, populations with opposing prognostic associations defined a markedly different survival pattern. *CXCL13+* T-cell-rich tumours with low *IGFBP5+* fibroblast abundance showed the most favourable outcomes, whereas high *IGFBP5+* fibroblast abundance was associated with shorter survival irrespective of *CXCL13+* T-cell abundance (HR = 2.1, p = 0.007) (**Figure 3F, Supplementary Table S9**). Thus, high *IGFBP5+* fibroblast abundance was associated with shorter survival even in patients with high tumour-proximal *CXCL13+* T-cell abundance.

Together, these analyses identified correlated groups of tumour-proximal cell populations associated with survival. Multiple longer-survival-associated immune populations formed a T-cell-enriched cluster with overlapping survival associations, whereas mesothelial cells and *IGHM+* B-plasma cells formed a distinct longer-survival-associated cluster. In contrast, *IGFBP5+* and *EGR1+* fibroblasts were associated with shorter survival but were not correlated with each other, with the association of *IGFBP5+* fibroblasts retained across a broader range of pairwise models.

### Tumour-intrinsic gene expression and neighbouring microenvironmental features are jointly associated with overall survival

Having identified tumour-intrinsic gene expression and tumour-proximal cell populations associated with overall survival, we next examined whether these features were related to one another and whether their combined associations provided additional information about patient outcome.

We first assessed correlations between tumour-intrinsic expression of survival-associated genes and the abundance of tumour-proximal cell populations across spatial distances. Multiple associations between gene expression and cell abundance were observed, with correlations generally strongest within 10–50 μm of tumour cells (**Figure 4A**, **Supplementary Table S10**). Higher expression of genes associated with longer survival correlated positively with the abundance of several immune populations also associated with longer survival. Most notably, *CHI3L1* expression correlated positively with the abundance of *CCL5+*, *IL32+*, *CXCL13+*, *CXCR4+* and regulatory T cells, as well as *CXCL9+* macrophages and *IGHG+* B-plasma cells. Similarly, *PTN* expression correlated positively with *CCL5+* T cell abundance, *LTF* expression with *IGHM+* B-plasma cell abundance, *CD44* expression with *CXCR4+* and *IL32+* T cell abundance, and *TNFRSF11B* expression with *CXCL9+* macrophage abundance. Conversely, expression of shorter-survival-associated genes, including *RNF43*, *PROX1*, *CTPS2*, *CCND1* and *ADA*, showed negative correlations with the abundance of mesothelial cells, while *COL4A5* expression was positively correlated with abundance of *IGFBP5+* fibroblasts.

**Figure 4.**
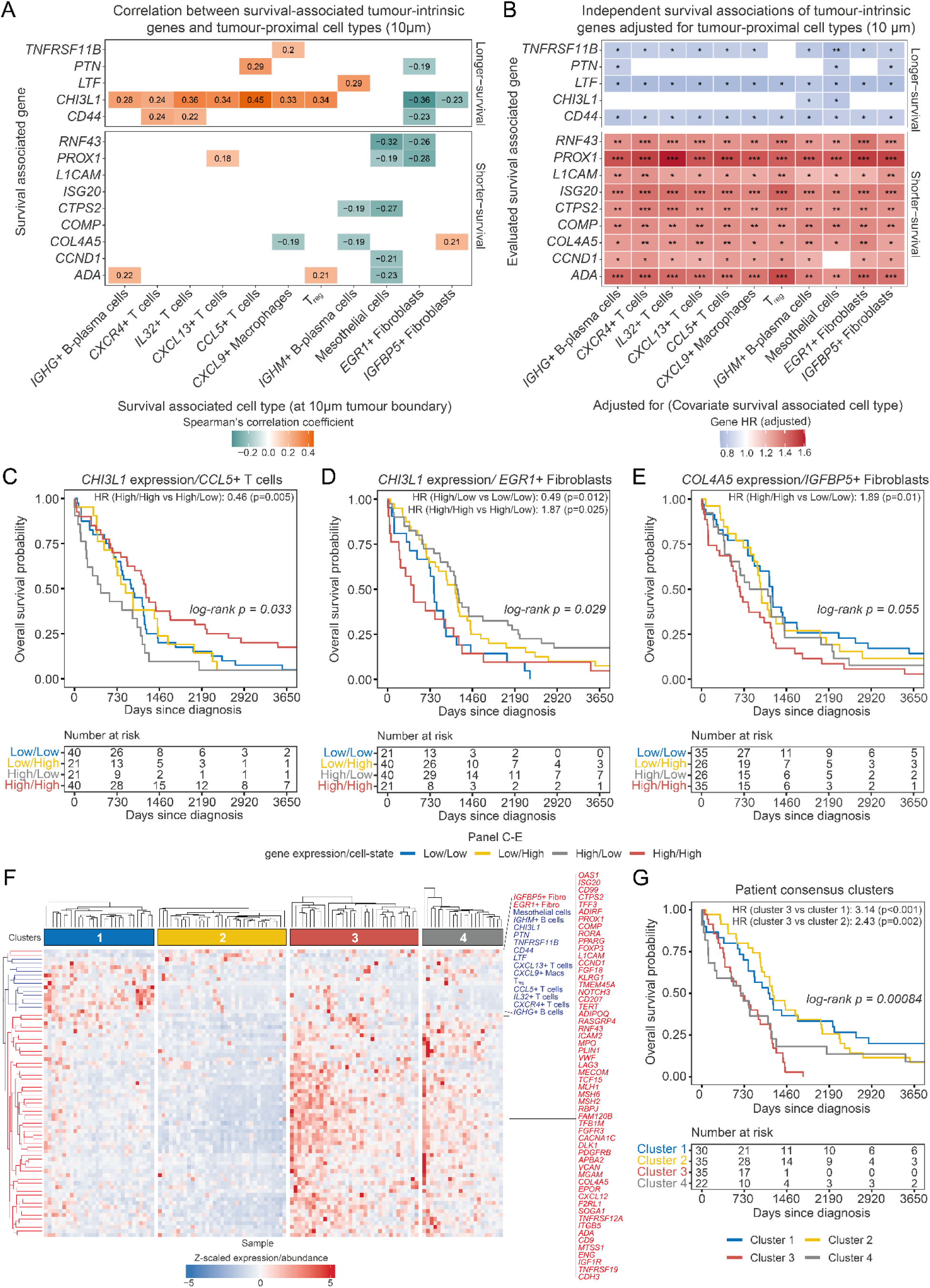
Tumour-intrinsic molecular features and tumour-proximal cellular populations jointly define prognostic tumour–microenvironmental states. **A.** Heatmap showing Spearman correlations between survival-associated tumour-intrinsic gene expression and the abundance of survival-associated cell populations within 10 μm of tumour cells. Only the statistically significant (p < 0.05) correlations are presented. **B.** Heatmap showing hazard ratios from pairwise joint Cox proportional hazards regression models evaluating each survival-associated tumour-intrinsic gene after adjustment for the corresponding tumour-proximal cell population. **C–E.** Kaplan–Meier analyses of overall survival for representative combinations of tumour-intrinsic gene expression and 10 μm tumour-proximal cell abundance. Patients were stratified by median expression and cell abundance into four groups: high gene/high cell, high gene/low cell, low gene/high cell and low gene/low cell. C. *CHI3L1* expression and *CCL5+* T-cell abundance, representing a favourable tumour–immune combination. D. *CHI3L1* expression and *EGR1+* fibroblast abundance, representing an opposing tumour–stromal combination. E. *COL4A5* expression and *IGFBP5+* fibroblast abundance, representing an adverse tumour–stromal combination. **F.** Heatmap showing z-scaled expression/abundance of survival-associated tumour-intrinsic genes and 10 μm tumour-proximal cell types across patients, with patients grouped according to consensus clustering (K = 4). Features are coloured in blue and red to indicate longer- or shorter-survival associations, respectively. The terminal edges and parent vertices of the row dendrogram are similarly coloured in blue and red to highlight the corresponding longer- and shorter-survival-associated features. **G.** Kaplan–Meier curves showing overall survival across the patient consensus clusters (K = 4). Kaplan–Meier curves displaying the first 10 years of overall survival; all analyses used the full survival duration. Hazard ratios and significance were estimated using Cox proportional hazards regression, with nominal p-values derived from the Wald statistic (*p < 0.05, **p < 0.01, ***p < 0.001). Global p-values for Kaplan–Meier survival curves were determined using the non-parametric log-rank test, which compares the survival distributions across patient groups over the observed follow-up period. For C–E, only significant contrasts are shown (p < 0.05). All reported p-values are nominal and were not adjusted for multiple testing.

To examine these relationships more comprehensively, we calculated pairwise Spearman correlations across all 61 survival-associated tumour-intrinsic genes and the 11 survival-associated tumour-proximal (10 μm) cell populations (**Figure S16**). Hierarchical clustering of the resulting correlation matrix revealed two broad groups of genes associated with shorter survival, together with a group comprising genes associated with longer survival and the previously identified T-cell-enriched immune populations. *IGHM+* B-plasma cells and mesothelial cells formed a separate correlated group, distinct from the T-cell-enriched populations. These findings indicate that the survival-associated features were not distributed independently across patients but exhibited broader patterns of correlated tumour-intrinsic expression and microenvironmental abundance.

We next asked whether tumour-intrinsic gene expression and neighbouring cell abundance within 10 μm of tumour cells retained their associations with survival after mutual adjustment or captured overlapping survival-associated information. Pairwise joint Cox regression models incorporating both tumour-intrinsic gene expression and neighbouring cell abundance showed that the associations of *CHI3L1* and *PTN* expression with longer survival were substantially attenuated after adjustment for several neighbouring immune populations that were also associated with longer survival including *CCL5+*, *CXCL13+*, *IL32+*, *CXCR4+* and regulatory T cells and *CXCL9+* macrophages. Conversely, the associations of these immune populations were similarly attenuated after adjustment for *CHI3L1* or *PTN* expression (**Figure 4B; Figure S17**). In contrast, *IGHM+* B-plasma cells and mesothelial cells retained associations with longer survival after adjustment for *CHI3L1* or *PTN*. These findings suggest that the survival associations of *CHI3L1* and *PTN* expression overlap with those of several neighbouring immune populations, whereas the associations of *IGHM+* B-plasma cells and mesothelial cells were largely retained after adjustment for either gene. Similarly, *COL4A5* expression retained its association with shorter survival after adjustment for *IGFBP5+* fibroblast abundance, whereas the association of *IGFBP5+* fibroblasts was attenuated after adjustment for *COL4A5*, consistent with overlap in their survival-associated information.

We extended these analyses by fitting pairwise Cox models across all survival-associated genes and cell populations (10 μm), adjusting the association of each feature with survival for every other feature in turn (**Figure S18**). The resulting matrix revealed substantial overlap in survival-associated information within the groups identified by correlation analysis. Many genes associated with shorter survival lost nominal significance after adjustment for other genes within the same correlated groups. Similarly, the survival associations of several T-cell-enriched populations were attenuated after adjustment for other populations within that group. Notably, the associations of *IGHM+* B-plasma cells and mesothelial cells were no longer significant after adjustment for one another, consistent with their correlated abundance and overlapping survival-associated information. Together, these analyses suggest that multiple individual gene and cell-population associations reflect broader, correlated patterns of tumour-intrinsic expression and microenvironmental composition.

We next asked whether combinations of tumour-intrinsic gene expression and tumour-proximal cell abundance could further stratify patient outcomes. Patients were stratified by median tumour-intrinsic gene expression and median cell abundance within 10 μm of tumour cells into four groups (**Figure 4C–E; Figure S19**). Representative gene–cell combinations were selected to illustrate different patterns of joint survival association. For *CHI3L1* and *CCL5+* T-cells, patients with high *CHI3L1* expression and high *CCL5+* T-cell abundance had longer survival than patients with high *CHI3L1* expression but low *CCL5+* T-cell abundance (HR = 0.46, p = 0.005; **Figure 4C**), demonstrating that *CCL5+* T-cell abundance further stratified outcomes among patients with high *CHI3L1* expression. For the combination of *CHI3L1* and *EGR1+* fibroblasts, patients with high *CHI3L1* expression and low *EGR1+* fibroblast abundance had longer survival than those with low *CHI3L1* expression and low *EGR1+* fibroblast abundance (HR = 0.49, p = 0.012). Among patients with high *CHI3L1* expression those with high *EGR1+* fibroblast abundance had shorter survival than those with low *EGR1+* fibroblast abundance (HR = 1.87, p = 0.025; **Figure 4D**), indicating the association between higher tumour-intrinsic *CHI3L1* expression and longer survival was attenuated in patients with high tumour-proximal *EGR1+* fibroblast abundance. Finally, patients with high *COL4A5* expression and high *IGFBP5+* fibroblast abundance had shorter survival than those with low *COL4A5* expression and low *IGFBP5+* fibroblast abundance (HR = 1.89, p = 0.01; **Figure 4E**), consistent with a combined adverse tumour-intrinsic and stromal state. Together, these analyses show that survival associations involving tumour-intrinsic gene expression can differ according to the abundance of specific neighbouring immune and stromal populations.

To determine whether the broader patterns of tumour-intrinsic expression and microenvironmental abundance identified above could distinguish patient groups, we performed consensus clustering using all 61 survival-associated tumour-intrinsic genes and the 11 survival-associated tumour-proximal (10 μm) cell populations (**Figure S20**). This identified four patient clusters with distinct combinations of gene-expression and cellular-abundance features (**Figure 4F**).

Cluster 1 was characterised by higher expression of genes associated with longer survival, greater abundance of the T-cell-enriched immune populations and relatively low expression of genes associated with shorter survival. Cluster 2 showed comparatively limited enrichment of either the longer- or shorter-survival gene-expression groups and the T-cell-enriched populations, but exhibited a stromal-cell-associated signal. Cluster 3 was characterised by higher expression of genes associated with shorter survival, whereas cluster 4 combined higher expression of both shorter- and longer-survival genes with greater fibroblast abundance.

Kaplan–Meier analysis revealed differences in overall survival across the four patient clusters (**Figure 4G**). Clusters 1 and 2 exhibited longer survival, whereas clusters 3 and 4 exhibited shorter survival, with cluster 3 showing significantly shorter survival than clusters 1 and 2 (HR = 2.43, p = 0.0022; HR = 3.14, p < 0.001, respectively). Although the clusters differed in their combinations of tumour-intrinsic expression and microenvironmental features, no significant differences were observed between clusters 1 and 2 or between clusters 3 and 4. Notably, clusters 3 and 4 both exhibited shorter survival despite differing in their expression of genes individually associated with longer survival and their fibroblast abundance. These findings indicate that distinct combinations of tumour-intrinsic gene expression and microenvironmental composition can be associated with similar survival outcomes.

The observed associations between tumour-intrinsic gene expression and neighbouring cell populations raised the possibility that tumour-derived signalling contributes to differences in the local microenvironment. To identify candidate signalling molecules among the survival-associated tumour genes, we cross-referenced these genes against ConnectomeDB2025^29^, identifying multiple genes encoding cell-surface or secreted signalling molecules. These included shorter-survival-associated genes (*CXCL12*, *KLRG1*, *VWF*, *DLK1*, *CDH3*, *ICAM2*, *COMP*, *ADA*, *ADIPOQ*, *FGF18*, *L1CAM*, *VCAN*, *COL4A5* and *CD99*) and longer-survival-associated genes (*CD44*, *LTF* and *PTN*) (**Supplementary Table S6**). Although *CHI3L1* was not annotated as a ligand in ConnectomeDB2025, it is also a secreted signalling protein, raising the possibility that tumour-derived *CHI3L1* contributes to interactions with neighbouring immune and stromal cells.

Together, these analyses show that survival-associated tumour-intrinsic gene expression and tumour-proximal cellular abundance exhibit correlated patterns across patients, with substantial overlap in their individual survival associations. The association between tumour-intrinsic gene expression and overall survival can also differ according to the abundance of specific neighbouring immune and stromal populations. Integrating these features identified four patient groups with distinct combinations of tumour-intrinsic expression and microenvironmental abundance, broadly separating patients with longer and shorter overall survival.

## Discussion

With the exception of near-ubiquitous *TP53* alteration, HGSOC is genomically heterogeneous, with few recurrent driver alterations present across the majority of tumours. Gene-expression profiling has nevertheless identified molecular subtypes and individual genes associated with clinical outcome, while immunohistochemical and transcriptomic studies have linked the abundance of multiple immune and stromal populations to shorter or longer survival^1,23–26^. However, bulk transcriptomic measurements inherently combine expression from malignant, immune and stromal cells, making it difficult to determine the cellular origin of survival-associated gene-expression signals. Similarly, estimates of microenvironmental composition derived from bulk tissue reflect the overall cellular content of the sampled region, are influenced by tumour fraction, and do not directly capture the abundance of specific cell populations in the immediate vicinity of tumour cells. Here, we applied Xenium spatial transcriptomics to treatment-naïve HGSOC tumours from 122 patients to address both limitations simultaneously. By analysing gene expression within tumour cells and quantifying the abundance of neighbouring immune and stromal populations, we identified tumour-intrinsic and microenvironmental features associated with overall survival and examined the relationships between them. Direct comparison with whole-tissue pseudobulk expression further enabled us to assess how resolving tumour-cell expression affected the identification of survival-associated genes.

At a nominal significance threshold, tumour-intrinsic expression of 61 genes and tumour-proximal abundance of 13 cell populations were associated with overall survival. Of the 61 genes, 18 showed concordant associations with overall survival (p < 0.05, same direction of effect) in at least one of three independent bulk transcriptomic datasets: TCGA-OV^23^, Tothill et al.^1^ and the 3,561-patient OTTA-SPOT cohort^24^ (**Supplementary Table S4**). Associations for 10 of these genes remained significant after Benjamini-Hochberg correction for multiple testing. Seven genes had published immunohistochemical evidence linking tumour-cell protein expression to overall survival in the same direction, including three also supported by the bulk transcriptomic datasets. Thus, 22 of the 61 tumour-intrinsic survival-associated genes had independent evidence of concordant survival associations at the RNA or protein level. For the remaining 39 genes, we did not identify independent evidence of concordant survival associations in HGSOC. Our direct comparison of tumour-cell and whole-tissue pseudobulk expression provides support for one potential explanation for this limited overlap with previous bulk studies. Of the 61 genes associated with survival using tumour-cell expression, only 15 were also identified using whole-tissue pseudobulk, while 46 did not reach the nominal significance threshold in the whole-tissue analysis. Conversely, 21 genes were associated with survival only using whole-tissue pseudobulk, and these predominantly reflected expression by non-malignant populations, particularly fibroblasts, but also macrophages, myofibroblasts, plasmacytoid dendritic cells and *IGHM+* B/plasma cells (**Figure S3**). These findings demonstrate directly within the same patients that survival associations can differ substantially depending on whether expression is measured specifically within malignant cells or across the complete cellular mixture. The mixed expression signal in whole-tissue profiles reflects both cell-type-specific expression and variation in cellular composition, which can obscure tumour-cell-specific survival associations while introducing associations arising predominantly from microenvironmental populations.

Several of the shorter-survival-associated genes without independent survival evidence in HGSOC have established roles in biological processes relevant to tumour proliferation and chemotherapy response. These included genes involved in DNA mismatch repair (*MSH2, MSH6*), cell-cycle progression (*CCND1*) and telomere maintenance (*TERT*). Notably, increased *IGF1R* expression and signalling have been experimentally linked to acquired cisplatin resistance in ovarian cancer cells, providing a potential mechanistic explanation for its association with shorter survival in our cohort^30,31^. Together, these findings extend the range of tumour-cell-intrinsic genes associated with HGSOC survival to include additional genes implicated in proliferation, genome maintenance and chemotherapy resistance.

Of the five genes associated with longer survival, *CD44* had independent concordant survival evidence in HGSOC^32^, whereas *CHI3L1*, *LTF*, *TNFRSF11B* and *PTN* did not. Higher expression of these four genes has been associated with longer survival in other cancers, including neuroblastoma, breast, renal, oral and lung cancers^33–40^. However, higher expression of *TNFRSF11B*, *CHI3L1* and *PTN* has been associated with shorter survival in other tumour types^41–44^, including a report linking higher *CHI3L1* expression to shorter survival in ovarian cancer^44^. These contrasting findings highlight the context-dependent prognostic associations of these genes and suggest that tumour-cell-resolved expression profiling may provide additional insight into survival associations that are not consistently observed in bulk transcriptomic studies.

Resolving the cellular origin of these associations was particularly important for genes expressed across multiple cell types. Several genes whose expression within malignant cells was associated with survival are more commonly recognised as markers of non-malignant populations. These included *FOXP3*, *ISG20* and *LAG3*, which are expressed by T-cell populations, and *VWF* and *ENG*, which are expressed by endothelial cells. Direct inspection of Xenium transcripts confirmed expression within annotated tumour-cell segments (**Figure S6**), while independent single-cell RNA-seq datasets and published tumour immunostaining provided additional evidence that these genes can be expressed by malignant cells in a subset of HGSOC tumours (**Figure S7-7**). Importantly, the association between gene expression and survival could differ according to the cellular compartment in which the gene was expressed. Most notably, higher *FOXP3* expression within malignant cells was associated with shorter survival, whereas greater tumour-proximal abundance of *FOXP3+* regulatory T cells was associated with longer survival. Together with the differences observed between tumour-cell and whole-tissue pseudobulk analyses, this illustrates how the survival association attributed to a gene can depend on both its expression within individual cell types and the relative abundance of those cell types within the sampled tissue. This distinction may help explain discrepancies between tumour-intrinsic and bulk expression analyses, although the basis for discordant associations involving other genes, including

*TFF3*, *MECOM* and *CTPS2*, remains unclear.

Beyond the individual survival associations, Elastic-Net modelling prioritised 14 tumour-intrinsic expression features from the 61 candidate genes. Among these, *PTN*, *LTF*, *CD44*, *TNFRSF11B*, and *CHI3L1* were associated with longer survival. Tumour-intrinsic expression of these genes was frequently associated with greater abundance of tumour-proximal immune populations that were themselves associated with longer survival. Several have been implicated in recruiting or stimulating pro-inflammatory immune populations or have been associated with immune-cell-rich microenvironments^45–52^. However, studies in other tumour types have also linked some of these genes to tumour-promoting functions^53–56^, indicating that their association with longer survival in HGSOC should not be interpreted as evidence of uniformly anti-tumour activity. In contrast, nine prioritised genes, *COMP*, *COL4A5*, *CCND1*, *CTPS2*, *PROX1*, *L1CAM*, *ISG20*, *ADA*, and *RNF43*, were associated with shorter survival. These genes did not converge on a single pathway but instead represented diverse biological functions^57–65^. Gene sets derived from the prioritised shorter- and longer-survival features further stratified survival within the cohort, indicating that combinations of tumour-intrinsic expression features can provide additional prognostic information beyond individual gene associations.

Our spatial neighbourhood analysis further showed that associations between microenvironmental cell populations and survival varied with their proximity to tumour cells, rather than simply reflecting their overall abundance. Immune populations associated with longer survival, including B-plasma cells, multiple T-cell populations and *CXCL9+* macrophages, generally showed progressively weaker survival associations as distance from tumour cells increased. While favourable associations involving broader T-cell, B-cell and macrophage infiltration have been reported previously, including studies examining tumour-centric spatial organisation^6,66–71^, the cellular resolution achieved here identified specific immune cell states, including *IL32+* T cells, *CXCR4+* T cells and *CXCL9+* macrophages, whose tumour-proximal abundance has not, to our knowledge, previously been associated with overall survival in HGSOC. Dendritic cells differed from these populations, with their association with longer survival evident predominantly at greater distances from tumour cells^72,73^. Published transcriptomic and immunohistochemical studies also provided concordant survival associations for several of the identified immune and stromal populations, including *IGHG+* plasma cells, *CXCL13+* T cells and regulatory T cells (**Supplementary Table S12**).

Correlation analysis revealed that tumour-intrinsic genes associated with longer survival clustered with several tumour-proximal T-cell populations and *CXCL9+* macrophages, whereas genes associated with shorter survival formed two broader groups (**Figure S16**). The co-occurrence of longer-survival-associated tumour-intrinsic expression and immune-cell abundance may reflect shared biological processes or interactions between tumour cells and neighbouring immune populations. For instance, *CXCL13+* T cells may contribute to B-cell recruitment and lymphoid organisation^74^, while *CCL5* and *CXCL9* signalling may contribute to the recruitment and organisation of neighbouring immune populations. Tumour-cell-derived *CCL5* has been shown to promote *CXCL9* expression in tumour-associated macrophages^75^, and *CXCL9*-mediated signalling can promote recruitment of tumour-reactive *CD8+* T cells ^75,76^. The co-occurrence of *CXCL13+* and *CCL5+* T cells, *CXCL9+* macrophages and B/plasma-cell populations in our cohort is consistent with these populations participating in overlapping immune-rich microenvironments, although the direction of signalling between them cannot be established from our data. These could include TLS-like niches, given the co-occurrence of T-cell and B/plasma-cell populations, although direct assessment of TLS organisation was beyond the scope of our study^70,77^.

A second, distinct pattern of correlated abundance involved *IGHM+* B/plasma cells and mesothelial cells, both of which were associated with longer survival (**Figure S15**). Their survival associations were no longer significant after adjustment for one another, suggesting that these populations capture overlapping prognostic information. Mesothelial cells maintained an association with longer survival across all distances, raising the possibility that mesothelial barriers or signalling contribute to differences in peritoneal tumour spread or microenvironmental organisation^78^. Whether their association with *IGHM+* B/plasma cells reflects a shared tissue environment, common underlying biological processes or direct interactions between these populations remains to be determined.

In contrast to these immune-rich microenvironments, both *IGFBP5+* and *EGR1+* fibroblasts were associated with shorter overall survival. Although stromal *EGR1* expression has previously been associated with poor prognosis in epithelial ovarian cancer, with EGR1 protein localised to α-SMA-positive stromal fibroblasts^79^, our spatial analysis localises this prognostic association to an *EGR1+* fibroblast population and further demonstrates that its survival association varies with proximity to tumour cells. Similarly, an *IGFBP5*-associated fibroblast signature has been associated with poor outcome in non-small-cell lung cancer^80^. Although fibroblast abundance has previously been associated with adverse outcomes in HGSOC, these findings resolve this association to specific fibroblast states that, to our knowledge, have not previously been linked to overall survival. Their abundances were not significantly correlated, suggesting that they represent distinct fibroblast-associated microenvironments rather than a single fibroblast-rich phenotype^81–83^. More broadly, CAFs in ovarian cancer can shape the tumour immune microenvironment through interactions with immune cells, extracellular-matrix remodelling and secretion of immunoregulatory factors^9,82,84–86^, with specific CAF states implicated in immune exclusion and immunosuppression^87^. The inverse relationships between fibroblast populations and several favourable tumour-proximal immune populations observed here may therefore reflect stromal microenvironments that are less permissive to local immune infiltration. Combined analyses were consistent with previous reports of survival differing according to the relative abundance of tumour-proximal immune and fibroblast populations^66,68^.

Pairwise Cox regression demonstrated substantial overlap in survival-associated information within the correlated groups of tumour-intrinsic genes and neighbouring cell populations, with many individual associations attenuated after adjustment for other features within the same group (**Figure S18**). Nevertheless, stratification by individual gene–cell combinations revealed differences in survival according to the surrounding cellular context. For example, higher tumour-intrinsic *CHI3L1* expression was associated with longer survival particularly when tumour-proximal *CCL5+* T cells were abundant, whereas this favourable association was attenuated in patients with high *EGR1*+ fibroblast abundance. Similarly, concurrent enrichment of shorter-survival-associated tumour-intrinsic gene expression and fibroblast populations was associated with shorter survival. These findings indicate that individual gene-expression associations can capture overlapping prognostic information with neighbouring cell populations, while their combined measurements can provide additional information about patient outcome.

Consensus clustering identified four patient groups with distinct combinations of tumour-intrinsic gene expression and microenvironmental composition. Two groups exhibited longer survival despite differing in their expression of genes associated with longer survival and abundance of T-cell-enriched immune populations. Similarly, two groups with shorter survival differed in their expression of longer-survival-associated genes and fibroblast abundance. Notably, one shorter-survival group exhibited higher expression of both longer- and shorter-survival-associated tumour-intrinsic genes together with greater fibroblast abundance, suggesting that expression of genes individually associated with longer survival does not necessarily correspond to favourable outcomes when accompanied by other adverse tumour-intrinsic or microenvironmental features.

The observed correlations between tumour-intrinsic gene expression and neighbouring cell abundance may arise through several non-exclusive mechanisms. Many of the genes identified here encode secreted ligands, extracellular matrix proteins or cell-surface molecules (**Supplementary Table S6**), raising the possibility that their expression by tumour cells influences neighbouring immune and stromal populations through paracrine, juxtacrine or extracellular matrix-mediated signalling^88^. Conversely, signals from immune or stromal cells neighbouring tumour cells may induce expression of survival-associated genes within tumour cells^89^, providing a potential mechanism through which the local microenvironment influences tumour-cell gene expression. The correlated enrichment of tumour-proximal immune populations in patients with longer survival may also reflect differences between patients in the effectiveness of anti-tumour immune surveillance. Previous studies of HGSOC long-term survivors have reported greater tumour immune infiltration together with genomic features associated with increased tumour immunogenicity, including homologous recombination deficiency and increased neoantigen burden^90,91^. Differences in immune surveillance may therefore reflect both tumour-intrinsic immunogenicity and host determinants of antigen presentation and immune recognition^92,93^. The observed associations with survival may also reflect intrinsic sensitivity or resistance to platinum-based chemotherapy^15^. The relative contributions of these mechanisms cannot be determined from our analyses. However, our previous observations that genetically distinct HGSOC subclones within the same tumour express distinct ligand repertoires and are associated with distinct local cellular environments are consistent with tumour-derived signalling contributing to microenvironmental differences, since these differences occur within the same patient and therefore cannot be explained solely by inter-patient variation in host immunity^5^.

Several limitations should be considered when interpreting these findings. The targeted 480-gene panel enabled cohort-scale spatial profiling but necessarily restricted the range of tumour and microenvironmental biology that could be examined. Approximately 220 genes were required to distinguish immune and stromal cell populations, limiting the number of genes available to interrogate tumour-cell expression. Consequently, many candidate survival-associated genes identified in previous bulk transcriptomic studies could not be included, including genes such as *CCNE1*^11^ that have previously been associated with survival in HGSOC^11^. Similarly, the restricted marker set may have prevented the identification or distinction of additional immune and stromal cell types and states associated with survival. Whole-transcriptome spatial profiling should enable more comprehensive identification of survival-associated tumour-cell gene expression and microenvironmental populations, as well as interrogation of ligand–receptor interactions and signalling relationships between tumour cells and neighbouring populations. Our survival analyses also did not adjust for BRCA1/2 mutation or homologous recombination deficiency (HRD) status, which is a major determinant of both clinical outcome and treatment selection in HGSOC, particularly in the maintenance poly(ADP-ribose) polymerase (PARP) inhibitor era^94^. Other determinants of outcome, including subsequent lines of therapy at relapse and secondary cytoreductive surgery, were not accounted for, and potential confounding by these factors cannot be excluded.

The breadth of features examined and the resulting multiple-testing burden require appropriate caution in interpreting individual associations. This challenge will be greater with whole-transcriptome spatial profiling, where substantially more genes, cell states and spatial relationships can be tested, increasing the need for larger cohorts and independent validation. Although a subset of the tumour-intrinsic and microenvironmental associations was supported by independent transcriptomic cohorts or immunostaining evidence, the spatially resolved associations and integrated analyses require independent validation. The prioritised gene features, pairwise Cox regression analyses and consensus clustering were all evaluated using features initially selected on the basis of their survival associations within the same cohort. Consequently, the resulting patient groups and combined survival associations should be considered exploratory. Furthermore, attenuation of individual survival associations after adjustment for correlated features does not establish biological redundancy or identify the mechanisms underlying their shared prognostic information. In addition, although tissue microarrays enabled spatial profiling across 122 patients, each core represents only a limited region of a tumour. Given our previous observation that tumour subclones within individual patients can occupy distinct microenvironments^5^, TMA-based sampling is likely to underrepresent the full extent of intratumoural spatial heterogeneity. Larger, independent cohorts incorporating broader spatial sampling, detailed genomic and clinical annotation, and whole-transcriptome measurements will therefore be important for determining the robustness and biological basis of the associations identified here. Applying these approaches to samples collected during or after therapeutic intervention may additionally reveal treatment-specific relationships between tumour-cell gene expression and the surrounding microenvironment.

In conclusion, cohort-scale Xenium profiling of HGSOC identified survival-associated gene expression within malignant cells and survival-associated immune and stromal populations surrounding those cells, including gene and cell-state associations not previously linked to HGSOC overall survival. Published transcriptomic and immunohistochemical studies provided independent support for a subset of these associations. Direct comparison with whole-tissue pseudobulk demonstrated that resolving expression specifically within tumour cells revealed survival associations not identified when expression was aggregated across the complete cellular mixture, while many whole-tissue-specific associations reflected expression from non-malignant populations. Integrating tumour-intrinsic gene expression with tumour-proximal cellular abundance revealed correlated groups of genes and cell populations with overlapping survival associations and identified distinct patient groups characterised by different combinations of tumour-intrinsic and microenvironmental features. These findings demonstrate that survival-associated tumour-cell gene expression and microenvironmental composition are interconnected, providing a basis for investigating the biological mechanisms underlying their relationships with patient outcome.

## Methods

### HGSOC patient cohort

This retrospective study included treatment-naïve primary HGSOC samples (n = 122) obtained during primary debulking surgery from patients diagnosed between 1986 and 2002, with clinical follow-up extending through August 2020. Primary tumours were localised to the ovary (n = 118) and fallopian tube (n = 4), and all tissues were histologically confirmed as high-grade (grades 2 and 3). Overall survival (OS) duration was calculated from the date of histopathological confirmation to the date of death from any cause, or censored at the date of last follow-up (**Supplementary Table S13**).

### HGSOC TMAs

Tissue microarrays (TMAs) with 1-mm cores were constructed using a Beecher tissue microarrayer using archival tumour blocks. Archival tumour samples used in the TMAs were obtained under Human Research Ethics Committee approval with a prospective consent for future ethically approved ovarian cancer research studies (SJGHC HREC Ref#085/#360). The ethics committee approved the use of these de-identified archival specimens in accordance with national ethical guidelines governing research using stored human tissue and health information. Follow-up data for all patients were obtained from hospital medical records and the hospital Cancer Registry.

### Custom 480-gene Panel design

A custom 480-gene panel was designed by integrating gene lists relevant to HGSOC from 14 major sources, along with manually selected genes (**Supplementary Table S2**). Genes were prioritised based on their recurrence across multiple lists, and a primary source was assigned to each gene. This included 187 genes from the PrOTYPE study^2^, 102 genes recurrently amplified or deleted in the TCGA-OV dataset^23^, 61 genes associated with myeloid and T/NK populations in HGSOC from Vázquez-García et al.^13^, 20 ligands and receptors expressed in tumour cells from ConnectomeDB2025^29^, 14 tumour and stromal genes from Schwede et al.^4^, 6 genes identified in the HGSOC plasma proteome from Álvez et al.^18^, 5 T cell subset marker genes from Yeh et al.^20^, 4 mismatch repair genes^19^ and 26 additional genes manually selected from literature searches and in house datasets.

### Xenium profiling

Formalin-fixed paraffin-embedded (FFPE) TMA blocks were rehydrated in an ice bath for 1 hour prior to sectioning. Sections of 5-μm thickness were prepared using a Leica microtome. To access deeper regions of each TMA block, where tumour cores likely exhibit better RNA integrity, the first 5–6 sections were collected onto standard Superfrost slides (Cat# 12-550-15, Fisher Scientific) and retained, and the subsequent section was reserved for the Xenium workflow. To accommodate the smaller capture area of Xenium slides, each full TMA section was divided into 2–3 smaller pieces using a surgical scalpel before floating in a 37 °C water bath. These sections were then mounted onto Xenium slides (PN-3000941, 10x Genomics). Slides were air-dried overnight at room temperature on a drying rack without a fan, followed by drying at 42 °C for 3 hours the next morning. Slides were stored in a desiccator at room temperature according to the manufacturer’s instructions (CG000578).

Spatial transcriptomic analysis was performed using the Xenium In Situ Gene Expression workflow with Cell Segmentation Staining, employing a custom 480-gene panel as described above. Xenium Slides & Sample Prep Reagents (PN-1000460, 10x Genomics) and Xenium Cell Segmentation Staining Reagents (PN-1000661, 10x Genomics) were used. Deparaffinisation and decrosslinking followed the manufacturer’s protocol (CG000578). Probe hybridisation, ligation, and amplification were performed according to the user guide (CG000749), with hybridisation maintained for 20 hours across all slides. Cell segmentation staining was carried out overnight for 17 hours at 4 °C, followed by autofluorescence quenching and nuclear staining. During the Xenium Analyzer setup (CG000584), prepared slides were kept on ice and protected from light. Finally, slides were processed using Onboard Analysis Version 3.3.

### Xenium data processing and transcript reassignments

Outputs from the Xenium Onboard Analysis included processed images, multimodal cell segmentations, coordinates of each cell segment and transcript, and cell-by-gene count matrices. Coordinates of each TMA core were obtained by drawing a polygon around them in Xenium Explorer (v4.1.1)^95^ to assign the respective patient identifiers.

To reconstruct the profiled TMA sections, the raw spatial transcriptomics data, including cell-by-gene expression matrices, individual transcript coordinates, and cell boundary polygons, were first extracted using the “SpatialData” Python package (Spatialdata v0.2.5.post0, Spatialdata_io v0.1.5)^96^. Spatial coordinates and cell boundary geometries were transformed sequentially for each section. An offset was applied to each section within a TMA to prevent spatial overlap while preserving the relative spatial arrangement within each section. Cell-by-gene matrices were then merged into a unified AnnData object.

To address technical artefacts introduced by imperfect cell segmentation including transcript spillover, heterotypic gene admixture, and artificial cell states near cell borders, we applied the Missegmented Transcript Inference Correction (MisTIC) framework^97^. MisTIC uses a variational Bayesian framework to probabilistically model transcript assignments by jointly evaluating the localised spatial distribution of individual RNA molecules and the global transcriptional profiles of discrete cell populations, without requiring morphological re-segmentation of the tissue images.

Transcripts designated as “unassigned” by the Xenium Onboard workflow were pre-filtered, and cell segments devoid of any transcript assignments were removed. The MisTIC correction pipeline (“txmistic” package v0.0.2) was executed independently for each TMA to account for TMA-specific transcript assignment characteristics.. No prior cell grouping or clustering information was provided, allowing the algorithm to establish cell-group assignments natively. For the initialisation of the MisTIC object, default parameters were utilised for TMA6, while for TMA5, an optional Leiden resolution of 0.5 was applied to optimise initial cluster definitions.

The structured spatial datasets were patchified to prepare minibatches for Stochastic Gradient Descent (SGD). Model parameters were initialised and optimised via a variational training loop executed for up to 20 epochs, with early stopping triggered automatically upon structural parameter convergence. Transcript reassignment and removal thresholds were optimised using a two-parameter grid search. Reassignment probability thresholds were evaluated via a grid ranging from 0.1 to 0.5 (with increment steps of 0.1), while absolute transcript removal thresholds were scanned from 0.0 to 0.3 (with increment steps of 0.1). Under a “best” selection criteria framework, the optimal combination of thresholds was programmatically determined to maximise signal denoising while preserving feature retention. Transcripts displaying a high probability of misassignment under these optimised thresholds were reassigned to their true cell of origin, yielding a corrected cell-by-gene expression matrix for downstream analyses.

### Single-cell and core-level quality control filtering

Following transcript reassignment and expression count adjustment, quality control was performed independently for each TMA to filter out low-quality cells and TMA cores. To filter out low-quality cells, standard filtering was performed using the “Scanpy” package (v1.10.0)^98^; cells were retained if they possessed a minimum of 10 total transcript counts and at least one detected gene. For TMA5, the total transcript counts threshold was reduced to 5 to account for the lower transcript counts observed in this TMA.

To evaluate the structural integrity of individual tissue cores across the unified dataset, the distribution of high-quality cellular yields per core was profiled by grouping cells based on their unique core identifier. A core-level threshold was applied to exclude any TMA core yielding fewer than 100 total cells. The remaining high-quality cell profiles and well-represented tissue cores were subsetted into a finalised AnnData object for downstream analyses.

### Automated marker-guided annotation of cell types

QC-filtered expression data were processed and clustered independently for each individual TMA to account for any batch-specific technical variance. To prepare the quality-filtered expression data for downstream analysis, count matrices were normalised to the median total counts per cell and log-transformed with Scanpy. Principal Component Analysis (PCA) was performed, and the top 50 principal components were evaluated. A neighbourhood graph was subsequently constructed to achieve coarse clustering; for TMA6, the first 10 principal components under default neighbour settings were used, whereas for TMA5, the neighbourhood graph parameters were explicitly set to n_neighbors=50 and all 50 PCs. Coarse-resolution unsupervised clustering was then executed using the Leiden algorithm with a resolution of 0.1, and cluster-specific marker genes were evaluated to map these baseline partitions to broader malignant and non-malignant compartments.

To independently infer high-resolution, continuous cell type identity scores, automated marker-based annotation was performed using the “decoupleR” Python package (v2.1.3)^99^. The marker sets used for cell-type annotation comprised 227 genes for non-malignant cell types, of which 97 were used for fine-level annotations, and 184 genes for tumour cells, including three markers of proliferating tumour cells (**Supplementary Table S14**). Cell lineage specific scores were calculated across the unified dataset using Univariate Linear Modelling (ULM) across the unified dataset. Cells were assigned to their top-scoring lineage based on the maximum ULM estimation score.

Ambiguous cellular identities were identified by assessing concordance between the unsupervised Leiden clusters and supervised decoupleR annotations. Specifically, a cell-level tumour/non-tumour mismatch filter was applied to identify discordant assignments. Cells belonging to tumour-associated Leiden clusters were flagged if decoupleR assigned them to a non-tumour lineage; conversely, cells belonging to non-tumour Leiden clusters were flagged if decoupleR classified them as either “Non-proliferating tumour” or “Proliferating tumour”. Cells with discordant assignments were excluded from downstream analyses. Following broad lineage assignment and mismatch filtering, fine-grained cellular subtypes, including subpopulations of T cells, B/plasma cells, fibroblasts, endothelial cells and monocytes/macrophages, were resolved using lineage-specific marker sets and the same decoupleR ULM workflow.

To construct a batch-corrected spatial representation across the integrated cohort, the deep learning framework ResolVI (scvi-tools v1.3.3)^100^ was used. The raw transcript count matrices were configured as model inputs, with the final fine-grained annotations as reference structures. To account for variation between instrumentation runs, array origin was specified as a categorical batch covariate. The model was trained under a semi-supervised configuration for up to 200 epochs with automated early stopping. Following convergence, the batch-corrected shared latent representation was extracted and used as the input for t-distributed stochastic neighbour embedding (t-SNE) to visualise the distribution of annotated cell lineages across the integrated cohort.

### Construction of tumour cell pseudobulk transcriptome profiles

To facilitate sample-level comparative analysis of tumour-intrinsic gene expression across the cohort, sample-specific pseudobulk expression profiles were constructed from the annotated datasets. The raw transcript count matrix was extracted and coupled with the corresponding metadata, isolating cells broadly classified as “Tumour” (encompassing all “Tumour” and “Proliferating tumour” cell lineages) for each patient. To ensure adequate cellular representation, patients were retained only if their corresponding tissue cores yielded a minimum of 20 aggregated tumour cells. Samples failing to meet this baseline cellular threshold were excluded from subsequent expression modeling.

For each patient, raw transcript abundances across all gene features were aggregated by summation to generate bulk-like raw count profiles. To account for variations in cellular yield across individual tissue cores and patient samples, these summed expression vectors were converted to mean pseudobulk profiles by dividing the aggregate transcript counts of each gene feature by the total number of tumour cells within that patient. The resulting mean-count matrices were then used for downstream comparative analyses.

Following pseudobulk profile construction, expression profiles from the independent TMAs were integrated to facilitate cohort-wide downstream survival analyses. Mean abundances were log_2_(count + 1) transformed. To reduce batch-variations while preserving biological heterogeneity, batch effect correction was performed on the log-transformed expression matrix using the ComBat function from the “sva” (v3.54.0) Bioconductor package^27^, implemented with non-parametric empirical Bayes priors. Samples were explicitly matched to their corresponding TMA vector to ensure correct alignment of the batch vector. The resulting batch-adjusted log-expression matrices were utilised for unified cohort-level survival analyses.

### Estimating the abundances of tumour neighbouring non-tumour cell types

We utilised the spatial coordinates of cells to construct a neighbourhood graph based on the distance between two cells using the Squidpy (v1.6.1) package^28^, independently for each discrete tissue core across a range of radial proximity thresholds (10, 25, 50, 100, 250, and 500 μm). If the distance was shorter than the given radius, these two cells were defined as neighbours, and an edge connecting them was established in the neighbourhood graph. This approach allowed us to identify cells that were physically close within each individual tissue core. Self-loops were excluded so that only interactions between distinct cells were considered. Subsequently, we quantified spatial abundance of non-malignant populations strictly within the defined spatial neighbourhoods surrounding tumour cells.

For each tissue core at every radial threshold, a neighbour count matrix was generated by tallying the number of connections linking tumour cells to each distinct non-malignant cell category. To generate unified patient-level metrics for the cohort, these raw interaction counts were summed across all TMA cores from a patient. To facilitate comparison and account for baseline variations in tissue area, cellular yield, and tumour abundance across individual samples, the aggregated counts were divided by the total number of non-tumour cells neighbouring tumour cells for each patient at each spatial radius. The resulting values represented the relative fractions of each of the 49 non-tumour cell states within the defined spatial neighbourhoods surrounding tumour cells. These fractions were retained for downstream survival modelling.

## Association of tumour-intrinsic gene expression and tumour-surrounding neighbourhood composition with clinical outcomes

Batch-adjusted tumour cell pseudobulk expression data were utilised to identify genes associated with clinical outcomes. Univariate Cox proportional hazards models were fitted independently for each gene using cohort-wide z-score-transformed abundances as continuous variables within the “survival” (v3.8.3) R package framework^101^. Genes showing non-specific expression across the whole dataset (*CAPS* and *CFD*) were excluded from downstream survival modelling to reduce the influence of non-specific expression patterns. Forest plots summarising the univariate risk estimates were generated using the “forestmodel” (v0.6.2) R package^102^.

To identify non-redundant tumour-intrinsic features associated with survival , we applied Elastic Net regularised Cox proportional hazards regression via the “glmnet” (v4.1.10) R package^103^. This feature selection was performed on the batch-corrected, log-transformed expression profiles of the outcome-associated genes initially screened via univariate analyses. The mixing parameter was established at α=0.5 to balance structural sparsity with group-correlation effects. Optimal penalty constraints were defined via internal 10-fold cross-validation, selecting the conservative one-standard-error rule threshold (λ-1se) to minimise overfitting.

To evaluate the joint associations of dual-gene expression profiles with patient survival, cohorts were stratified into High or Low expression groups based on the median abundance of each gene independently. These classifications were subsequently integrated into a unified four-tier grouping scheme- High/High, High/Low, Low/High, and Low/Low. Survival probabilities across these strata were compared using Kaplan-Meier (KM) curves and associated global log-rank p-values generated via the “survminer” (v0.5.1) R package^104^. Pairwise Cox regression analyses were then performed to estimate hazard ratios for each combined expression stratum relative to the baseline Low/Low reference group.

An analogous survival analysis pipeline was implemented for tumour-neighbouring non-tumoural cell abundances. However, due to the sparse spatial distribution of certain cell states across individual tissue cores, we opted for a stratified quantile comparison. Patients were partitioned into quartiles based on the abundance distribution of each cell state across the evaluated radii. Cox regression modelling was then performed exclusively to compare the highest quartile (top 25%) directly against the baseline lowest quartile (bottom 25%). Combined KM-curves with associated global log-rank p-values were generated using the same methodology described above; patients were stratified into high and low groups via median splits for multiple cell states, which were subsequently integrated to evaluate the co-implications of multi-cell type spatial patterns.

Pairwise joint Cox proportional hazards models were used to evaluate whether prognostic associations among tumour–proximal features were retained after adjustment for each other. All features were z–score standardised across patients. For each pair, models of the form Survival ∼ feature A + feature B were fitted, enabling the association of each feature to be assessed while adjusting for the other. This generalised framework was applied to all combinations of survival–associated features, including cell–type, gene–expression, and mixed gene–cell–type pairs.

To investigate putative interactions between tumour-intrinsic gene expression and the cellular composition of the microenvironment, pairwise Spearman rank correlation coefficients were calculated between selected gene expression profiles and neighbouring cell type fractions across multiple radii. We then applied a combinatorial, stratified Cox regression approach to screen for joint survival associations. Specifically, survival outcomes were contrasted between patients presenting with high abundances of both the target gene and the neighbouring cell states against the baseline cohort exhibiting low abundances for both features. Similarly, survival trajectories were compared between the discordant strata (high gene/low cell state versus low gene/high cell state) to map out distinct microenvironmental survival risks.

To evaluate the collective transcriptional activity of our identified prognostic gene sets across individual patients, we performed single-sample Gene Set Enrichment Analysis (ssGSEA) using the GSVA (v2.0.7) R package^105^. Enrichment scores were calculated directly from the batch-adjusted, log_2_-transformed pseudo-bulk tumour expression matrix. We evaluated distinct gene signature sets (9-shorter-survival gene set and 5-longer-survival gene set) derived from the regularised candidate features prioritised via Elastic-Net analysis described above. The ssGSEA calculations were executed utilising the ssgseaParam configuration framework without inter-sample range normalisation, thereby preserving the raw, magnitude-dependent single-sample enrichment scores. Prior to downstream clinical correlation and multivariate survival modelling, all resulting continuous gene set enrichment metrics were z-score normalised across the patient cohort to ensure direct baseline comparison of clinical effect sizes.

To summarise the combined prognostic behaviour of the tumour–intrinsic gene expression patterns, we constructed a unified regularised risk score using the gene features retained by Elastic–Net analysis. Expression values for each selected gene were first z–score normalised across patients. For each individual, the unified score was obtained by weighting these normalised expression values by their corresponding Elastic–Net coefficients (β) and summing the resulting contributions to generate a single continuous index, which was subsequently used in the survival analyses.

To assess the out-of-sample prognostic performance of the tumour-intrinsic gene expression model, nested 10-fold cross-validation was performed using the complete 480-gene expression panel. Within each outer training fold, gene expression was standardised using training-set parameters, and genes associated with overall survival were selected by univariate Cox regression (nominal p < 0.05). The selected genes were used to fit the previously described Elastic Net Cox regression model, which was then applied to the held-out fold to generate out-of-fold risk scores. This process was repeated across all 10 folds, ensuring that each patient’s score was generated without using their survival outcome during model development.

To evaluate the longitudinal discriminative behaviour of the unified regularised risk score, time–dependent receiver operating characteristic (timeROC) curves were generated at 1–, 2–, and 5–year clinical timepoints using the timeROC (v0.4.1) R package^106^. The procedure accounts for censoring via inverse probability of censoring weighting (IPCW), and 95% confidence intervals were derived using influence–function–based variance estimation. False–positive and true–positive coordinates were extracted from the fitted models for visualisation.

### Evaluation of tumour-intrinsic prognostic signals in independent bulk transcriptomic datasets

To assess whether tumour-cell-intrinsic survival-associated signals identified by spatial profiling were recapitulated in conventional bulk transcriptomic data, we evaluated the 61 survival-associated genes and the derived shorter-survival and longer-survival gene-set scores across independent HGSOC bulk transcriptomic datasets. Tothill et al.^1^ microarray dataset (GSE9891) was obtained from the gene expression omnibus (GEO) using the GEOquery package^107^. Publicly available processed expression matrices were used, with expression values log2-transformed where required. Microarray probes were annotated to gene symbols using the corresponding platform annotation. Probes without an assigned gene symbol were excluded, and multiple probes mapping to the same gene were collapsed by taking the mean expression value. Corresponding clinical survival information was obtained from KMplotter^108^ and restricted to grade 2–3 serous ovarian cancer samples with available overall survival information. Separately, TCGA-OV^23^ RNA-seq data were obtained from the Genomic Data Commons (GDC) using TCGAbiolinks^109^. Primary tumour samples were selected, gene-level unstranded STAR counts were aggregated for duplicate gene symbols, TMM-normalised using tidybulk^110^, and log2-transformed. Clinical information was obtained from TCGA clinical data, and samples were restricted to grade 2–3 tumours with available overall survival information. For evaluation of individual survival-associated genes, expression values were z-score transformed independently within each dataset and assessed as continuous covariates in Cox proportional hazards regression models. Hazard ratios (HRs) were estimated from the Cox models, with statistical significance assessed using the Wald statistic.

### Analysis of independent HGSOC single-cell RNA-seq datasets

To assess the expression patterns of the 61 survival-associated genes identified by Xenium profiling in independent HGSOC single-cell RNA-sequencing datasets, we analysed publicly available datasets from MSK SPECTRUM (*n* = 41)^13^, Zheng et al. (*n* = 10)^111^ and Zhang et al. (*n* = 11)^15^. Raw count matrices were normalised to total library size and log-transformed using Scanpy. Cell-type annotations provided with each dataset were used to identify tumour cells, with cancer/epithelial cell populations mapped to a common tumour category. Expression of the 61 survival-associated genes was then examined across tumour cells at the individual sample/patient level alongside the annotated non-tumour cell populations.

### Consensus clustering of patients based on survival-associated tumour-intrinsic genes and tumour-proximal cell types

Survival-associated tumour-intrinsic gene expression and 10 μm tumour-proximal cell-type abundances were integrated into a single feature-by-patient matrix comprising 61 tumour-intrinsic genes and 11 tumour-proximal cell types. To place gene expression and cell-abundance features on a comparable scale, each feature was independently z-scaled across patients. Consensus clustering was performed using the ConsensusClusterPlus^112^ package with 10,000 resampling iterations. In each iteration, 80% of patients and 90% of features were randomly sampled, followed by partitioning around medoids (PAM) clustering using Spearman distance. Consensus matrices were generated for K values ranging from 2 to 7, and after assessing clustering stability across the different K values, K = 4 was selected for downstream analysis. Patient assignments from the four-cluster solution were used to visualise the integrated gene and cellular profiles and to evaluate differences in overall survival using Kaplan–Meier analysis and Cox proportional hazards regression.

## Data availability

The processed spatial transcriptomic data generated in this study are available as a consolidated AnnData object from Figshare (https://doi.org/10.6084/m9.figshare.34052868). The Xenium outputs are being deposited in the Gene Expression Omnibus (GEO) and will be made publicly available upon acceptance of the manuscript.

## Code availability

Code for the key analyses of the Xenium data and survival associations is available in our GitHub repository (https://github.com/Kamran-Khan96/Xenium_HGSOC_spatial). All other analyses were performed using standard tools cited in the manuscript.

## Authors contributions

ARRF conceived the project. MAAKK, WL, RH, YH, and PL performed the analyses. ARRF and ED designed and curated the custom Xenium panel. SB, YY and PAC were responsible for ethics, clinical annotation, and interpretation of the TMA samples. ARRF, PAC, WL, and MAAKK drafted the manuscript, with input and contributions from all authors.

## A.I. usage statement

ChatGPT was used to suggest changes in the manuscript that would improve flow and logic in the manuscript text. All analyses, interpretation of results, and scientific conclusions were performed and verified by the authors.

## Acknowledgements

We would like to acknowledge the patients who participated in this study. We would like to acknowledge Jane Thompson, Barry Iacopetta, Nik Zeps and Anne-Marie Shearwood who had the foresight to generate the TMAs used in this study in the early 2000s. We would also like to acknowledge Maria Beilin, Adeline Tan and Lisa Spalding for advice on the TMAs.

## Supplementary Materials

### Supplementary Tables

**Supplementary Table S1:** Clinicopathological features of the included HGSOC patients.

**Supplementary Table S2:** Target gene list and associated probe information for the custom 480-gene Xenium panel.

**Supplementary Table S3:** R^2^ values and associated p-values from pairwise linear regression analysis of median abundances of the 17 cell type lineages across Xenium slides.

**Supplementary Table S4:** Supporting evidence for tumour-intrinsic survival-associated genes across independent datasets. HRs, nominal P values and BH-adjusted P values are shown for the Xenium cohort and independent bulk transcriptomic datasets (OTTA-SPOT, TCGA-OV and Tothill et al.). Directional concordance with bulk datasets is summarised as the number of concordant datasets over those available for each gene. Previously reported intratumoural IHC evidence and HPA protein staining in serous ovarian carcinoma are shown where available. The final column reports the median percentage of tumour cells expressing each gene across samples in independent ovarian cancer scRNA-seq datasets, with the lowest–highest range in parentheses. HR >1 indicates shorter survival and HR <1 indicates longer survival. (NA, not available; BH, Benjamini–Hochberg; IHC, immunohistochemistry; HPA, Human Protein Atlas)

**Supplementary Table S5:** Survival associations of genes using tumour-cell only pseudo-bulk and whole-tissue pseudo-bulk expression. Hazard ratios (HRs) and nominal p-values from univariate Cox regression analyses are reported.

**Supplementary Table S6:** Copy-number alterations (CNAs) and ligand–receptor annotations of survival-associated genes in the TCGA ovarian cancer (OV) cohort. Frequencies of gene amplification and deletion, together with the corresponding chromosomal cytoband information, were retrieved from the cBioPortal database. Ligand–receptor classifications were obtained from ConnectomeDB2025^29^.

**Supplementary Table S7:** Coefficients (β) of the 61 outcome-associated genes after Elastic-net regularisation.

**Supplementary Table S8:** Pairwise joint Cox regression models evaluating whether prognostic associations of spatially defined tumour-proximal cell populations were retained after adjustment for each other across spatial radii.

**Supplementary Table S9:** Survival associations of tumour-proximal cellular population combinations defined by median-split abundance (High-High, High-Low, Low-High and Low-Low) across spatial radii. Cox regression hazard ratios and associated nominal p-values for each pairwise comparison are reported.

**Supplementary Table S10:** Correlations between tumour-intrinsic gene expression and tumour-neighbouring cell type abundances across incremental spatial radii.

**Supplementary Table S11:** Survival associations of tumour-intrinsic gene expression and tumour-proximal cell type abundance combinations defined by median splits (High gene–High cell type, High gene–Low cell type, Low gene–High cell type and Low gene–Low cell type) across spatial radii. Cox regression hazard ratios and associated nominal p-values for each pairwise comparison are reported.

**Supplementary Table S12:** Supporting evidence for tumour-proximal cell states across independent datasets. Summary of marker genes for each survival-associated tumour-proximal cell state identified in the Xenium cohort. HRs, nominal P values and BH-adjusted P values are shown for the OTTA-SPOT, TCGA-OV and Tothill et al. bulk transcriptomic datasets. Directional concordance with the Xenium-associated survival direction is summarised as the number of concordant datasets over those available for each gene. The final column lists published cell-level evidence supporting the corresponding cell state where available. HR <1 indicates longer survival and HR >1 indicates shorter survival. NA, not available; BH, Benjamini–Hochberg.

**Supplementary Table S13:** Clinical data of the cohort profiled in this study.

**Supplementary Table S14:** Gene markers used for the supervised lineage level and fine-grained cell subtype annotations.

### Supplementary Figures

**Figure S1**: **Overview of clinicopathological covariates, cell**–**type abundance distributions, and single**–**cell lineage resolution across the unified cohort**. **A.** Multivariate Cox proportional hazards forest plot evaluating the association of baseline clinicopathological features and continuous patient-level tumour cell abundance with patient survival outcomes. Data points represent hazard ratios (HR), horizontal error bars denote 95% confidence intervals (CI), and associated p-values are displayed on the right. **B.** Box-and-whisker plots depicting the cell lineage abundance distributions within the broad Tumour, Stromal, and Immune coarse-compartments across individual samples, sorted by median cell lineage abundance. The vertical axis is displayed on a pseudo-log scale. **C.** Scatter plot showing a linear regression model of the median abundances for all 17 broad cell types compared between the two profiled TMAs (TMA5 versus TMA6). The dashed line represents the identity line, and the solid orange line indicates the linear fit. **D.** t-SNE visualisations of the deep-learning-derived, integrated latent space distribution of the single cells colour-coded by individual technical Xenium runs; the right embedding displays a continuous colour gradient of patient-specific overall survival (OS) duration (days). **E.** Dot plot matrices detailing the high-resolution marker profiles used to sub-cluster broad lineages (B-plasma cells, T cells, Monocytes-Macrophages, Fibroblasts, and Endothelial cells) into fine-grained cellular states. The size of each dot represents the percentage of cells within a group expressing a given marker, and the colour scale reflects log-fold change compared to all other subtypes within that broad lineage. **F.** Box-and-whisker plots showing the percentage abundance distributions of the fine-grained cellular states within their respective compartments across the unified patient cohort. The vertical axis is displayed on a pseudo-log scale.

**Figure S2: Cross**–**TMA concordance of tumour**–**cell pseudobulk gene expression profiles. A.** Principal Component Analysis (PCA) plots of sample-specific tumour cell pseudobulk gene expression profiles across the two profiled TMAs (TMA5 (blue) and TMA6 (red)) before (left) and after (right) batch correction using ComBat. **B.** Correlation of median gene expression levels (log_2_ Count) for all 480 genes between the two TMAs. Association estimates (R^2^ and p-value) were derived from a linear regression model.

**Figure S3: Comparison of survival-associated genes identified using tumour-cell and whole-tissue pseudo-bulk expression. A.** Venn diagram showing the overlap between 61 survival-associated genes identified using tumour-cell pseudo-bulk expression and 36 genes identified using whole-tissue pseudo-bulk expression. **B.** Scatter plot comparing hazard ratios (HRs) for genes identified in the two analyses. **C.** Dot plot showing the expression of the 21 genes uniquely associated with survival in the whole-tissue pseudo-bulk analysis across 51 annotated cell types.

**Figure S4: Expression of survival-associated genes in tumour cells in the Xenium data.** Dot plot showing expression of the 61 tumour-intrinsic survival-associated genes within tumour cells across individual patients. Dot size indicates the proportion of tumour cells expressing each gene, and colour intensity indicates scaled mean expression within each patient. Gene order follows the order shown in Figure 2B.

**Figure S5: Expression of survival-associated genes across non-tumour cell populations in the Xenium data**. Dot plot showing expression of the 61 tumour-intrinsic survival-associated genes across the annotated 49 non-tumour cell populations. Dot size indicates the proportion of cells expressing each gene, and colour intensity indicates scaled mean expression within each displayed cell type. Gene order follows the order shown in Figure 2B.

**Figure S6: Spatial distribution of survival-associated gene transcripts across HGSOC tissue sections.** Representative Xenium images showing the spatial localisation of survival-associated gene transcripts (yellow dots) across the 61 survival-associated genes relative to tumour cells (red) and immune/stromal populations (grey).

**Figure S7: Protein expression of survival-associated genes in HGSOC.**

Representative Human Protein Atlas (HPA) immunohistochemistry images showing protein expression of survival-associated genes in serous cystadenocarcinoma ovarian cancer tissue. Only genes with detectable tumour cells immunostaining in the HPA are shown; the remaining survival-associated genes either exhibited no detectable staining or were not detected.

**Figure S8: Expression of survival-associated genes in independent HGSOC single-cell RNA-seq datasets.** Dot plots showing expression of the 61 tumour-intrinsic survival-associated genes identified in the Xenium cohort across three independent ovarian cancer single-cell RNA-seq datasets, shown in the following order: MSK SPECTRUM, Zhang et al., and Zheng et al. Tumour cells are displayed separately for each patient, whereas non-tumour populations are grouped according to their annotated cell type. Dot size indicates the proportion of cells expressing each gene, and colour intensity indicates scaled mean expression within each displayed group. Gene order is consistent across all three datasets and follows the order shown in Figure 2B.

**Figure S9: Heatmap displaying pairwise Spearman’s correlation coefficients between the 61 survival-associated genes.** Only statistically significant (p < 0.05) values are presented within the heatmap.

**Figure S10: Feature selection and coefficient penalisation via Elastic-Net regularised regression. A**. Cross-validation error curve showing partial likelihood deviance across a range of log-transformed tuning parameters (log(λ)). Red dots represent mean deviance across data folds, with error bars indicating ±1 standard error (SE). Vertical lines designate the minimum criteria threshold (log(λ_min_), grey dotted line) and the most parsimonious model threshold within 1-SE of the minimum (log(λ_1se_), red dashed line), and the associated number of retained prognostic features for each threshold. **B**. Coefficient trace paths for the 61 survival-associated genes across varying penalisation levels (log(λ)). Active features selected at the log(λ_1se_) boundary (red dashed line) are highlighted in colour, demonstrating the shrinkage of non-redundant features to zero. **C.** Heatmap displaying pairwise Spearman’s correlation coefficients between the 14 Elastic-net prioritised survival-associated genes. Only statistically significant (p < 0.05) values are presented within the heatmap. **D.** Heatmap showing hazard ratios (HRs) from pairwise joint Cox regression models of the 61 survival-associated genes. Each model evaluated the survival association of one gene after adjustment for a second gene. **E.** Multivariate Cox regression analysis of the 9-gene shorter survival and 5-gene longer survival gene set scores. Continuous set scores were z-score normalised across the cohort prior to model fitting. The normalised scores were then co-modelled simultaneously to determine their mutually adjusted prognostic associations. Significance metrics for the Cox proportional hazard models were derived using the Wald statistic.

**Figure S11: Prognostic evaluation and time**–**dependent validation of the unified regularised risk score. A.** Kaplan–Meier survival curve demonstrating patient stratification based on a median split of the continuous unified regularised risk score. Kaplan–Meier curves display only the first 10 years of overall survival for visualisation purposes; analyses used the full survival duration. Hazard ratio (HR) with corresponding 95% confidence intervals (CI), p-value from Wald statistic, and the log–rank p-value are indicated. **B.** Time–dependent receiver operating characteristic (timeROC) curves assessing the discriminative behaviour of the continuous unified score at 1–year (red), 2–year (blue), and 5–year (green) follow–up timepoints. Area under the curve (AUC) values and associated 95% CIs are reported for each interval. **C.** Kaplan–Meier survival curve based on patient-level out-of-fold unified risk scores generated using nested 10-fold cross-validation. In each of 10 rounds, 10% of patients were held out, while feature selection and Elastic-Net model fitting were performed using the remaining 90% of patients; the resulting model was then applied to the held-out patients to generate their out-of-fold scores. Patients were stratified by the median out-of-fold risk score. HR with corresponding 95% CI, p-value from Wald statistic, and log-rank p-value are indicated.

**Figure S12: Spatial distribution profiles and multi**–**radius survival analyses of tumour**–**adjacent microenvironmental components.** All spatial abundances represent the fraction of each tumour-proximal non–tumour cell type out of the total tumour-proximal non–tumour cell population within each patient, computed independently at each proximity radius. **A.** Stratified box plots showing the proportional abundance of non–malignant cell subtypes within the immediate, intermediate, and distant neighbourhoods of tumour cells across six predefined spatial radii (10, 25, 50, 100, 250, and 500 μm). **B.** Heatmap demonstrating hazard ratios (HR) derived from independent univariate Cox proportional hazards regression models evaluating continuous z–score–transformed spatial abundances across all spatial distances. **C.** Heatmap summarising categorical Cox regression HRs comparing extreme patient quartiles (75th percentile versus 25th percentile of abundances) for each non–malignant cell subtype (only cell types which were not presented in Figure 3B) across the six incremental proximity radii. In panels B–C, significance metrics for the Cox models were derived using the Wald statistic (*p < 0.05, **p < 0.01, ***p < 0.001).

**Figure S13: Univariate Cox proportional hazards regression evaluating the prognostic associations of cell**–**type abundances.** Abundances reflect the fraction of cell types out of total cells within each patient, across hierarchical cellular granularities. All cell–type fractions were transformed using z–score standardisation prior to modelling. **A.** Major cellular compartments, comprising tumour, stromal, and immune classes. **B.** Intermediate lineage–level cell types, including fibroblasts, endothelial cells, myeloid, lymphoid subsets, and additional stromal lineages. **C.** Fine–grained transcriptionally resolved subtypes and functional states, spanning fibroblast, monocyte–macrophage, endothelial, B–cell, and T–cell subsets. Data points represent hazard ratios (HR), with horizontal bars indicating 95% confidence intervals (CI). Associated p–values from Wald statistic are reported for each cell–type category.

**Figure S14: Spatial co**–**abundance correlation matrices of prognostically significant cell subtypes across incremental microenvironmental radii.** Heatmaps display Spearman’s rank correlation coefficients between the spatial abundances of non–tumour cell subtypes within matching spatial distances. For each panel, the matrix includes only those cell subtypes that demonstrated a significant survival association under extreme–quartile split comparisons at that specific radius. Only statistically significant (p < 0.05) correlation coefficient values are presented.

**Figure S15: Pairwise joint Cox regression of survival-associated tumour-proximal cell states at the 10 μm radius**. Heatmap showing hazard ratios from pairwise joint Cox regression models in which each survival-associated cell state was evaluated after accounting for a second cell state. Significance was assessed using the Wald statistic (*p < 0.05, **p < 0.01 and ***p < 0.001). Only statistically significant (p < 0.05) associations are shown; non-significant associations are shown as white tiles.

**Figure S16: Spearman correlation among survival-associated tumour-intrinsic genes and tumour-proximal cell populations (10 μm).** Heatmap showing pairwise Spearman correlations among the 61 survival-associated tumour-intrinsic genes and 11 tumour-proximal cell populations. Only statistically significant (p < 0.05) correlations are shown. Dendrograms show hierarchical clustering of the features based on their correlation profiles.

**Figure S17: Heatmap showing hazard ratios from joint Cox proportional hazards models evaluating each survival-associated tumour-proximal cell population at 10 μm after adjustment for the survival-associated tumour-intrinsic genes**. Significance was assessed using the Wald statistic (*p < 0.05, **p < 0.01 and ***p < 0.001). Only statistically significant (p < 0.05) associations are shown; non-significant associations are shown as white tiles.

**Figure S18: Heatmap showing hazard ratios from pairwise joint Cox proportional hazards regression models evaluating each of the 61 survival-associated tumour-intrinsic genes and 11 tumour-proximal cell populations after adjustment for each other feature individually.** Significance was assessed using the Wald statistic (*p < 0.05, **p < 0.01 and ***p < 0.001). Only statistically significant (p < 0.05) associations are shown. Dendrograms show hierarchical clustering of the features, with feature ordering consistent with Figure S16.

**Figure S19: Combined prognostic evaluations of tumour-intrinsic gene expression and tumour-proximal cell type abundances within 10 μm radius of tumour cells.** Patients were stratified by median splits for both tumour-intrinsic gene expression and neighbouring cell type abundance, generating four groups within each spatial niche: High gene–High cell type, High gene–Low cell type, Low gene–High cell type, and Low gene–Low cell type. **Left panel:** Heatmaps displaying hazard ratios (HRs) comparing High gene–High cell type versus Low gene–Low cell type, evaluating the combined prognostic association of high tumour-intrinsic gene expression and high neighbouring cell type abundance. **Middle panel**: Heatmaps displaying HRs comparing High gene–High cell type versus High gene–Low cell type, evaluating the association of neighbouring cell type abundance among patients with high tumour-intrinsic gene expression. **Right panel**: Heatmaps displaying HRs comparing High gene–High cell type versus Low gene–High cell type, evaluating the association of tumour-intrinsic gene expression among patients with high neighbouring cell type abundance. Significance was assessed using the Wald statistic. Only statistically significant (p < 0.05) HRs are shown. The x-axes include only cell types that demonstrated significant survival associations under extreme-quartile split comparisons at the 10 μm radius.

**Figure S20: Consensus clustering analysis of patient subgroups using 61 survival-associated tumour-intrinsic genes and 11 tumour-proximal (10 μm) cell-type abundances.** Summary of ConsensusClusterPlus outputs across K = 2–7, showing cluster stability and consensus structure across candidate numbers of patient clusters, with K = 4 selected for downstream analyses.

