## Supplementary Table S1 for "Spatial transcriptomics reveals tumour cell-intrinsic and microenvironmental features associated with survival in high-grade serous ovarian carcinoma"

**Supplementary Table S1: Clinicopathological features of the included HGSOC patients.**

| **Age at diagnosis** | Median 65 years (35 – 92 years) | |
| --- | --- | --- |
| **Overall survival duration** | Median 1006 days (0 – 8604 days) | |
| **Vital status** | Dead | 118 |
|  | Alive | 4 |
| **Residual disease** | R = 0 | 111 (91%) |
|  | R < 1 | 3 (2.46%) |
|  | R = 1-2 | 4 (3.28%) |
|  | R > 2 | 4 (3.28%) |
| **Site** | Ovary | 118 (96.7%) |
|  | Fallopian Tube | 4 (3.3%) |
